# CTCF is a Barrier for Totipotent-like Reprogramming

**DOI:** 10.1101/2020.12.20.423692

**Authors:** Teresa Olbrich, Maria Vega-Sendino, Desiree Tillo, Wei Wu, Nicholas Zolnerowich, Andy D. Tran, Catherine N. Domingo, Mariajose Franco, Marta Markiewicz-Potoczny, Gianluca Pegoraro, Peter C. FitzGerald, Michael J. Kruhlak, Eros Lazzerini-Denchi, Elphege P. Nora, Andre Nussenzweig, Sergio Ruiz

## Abstract

Totipotent cells have the ability of generating embryonic and extra-embryonic tissues^1,2^. Interestingly, a rare population of cells with totipotent-like potential was identified within ESC cultures^3^. These cells, known as 2 cell (2C)-like cells, arise from ESC and display similar features to those found in the totipotent 2 cell embryo^2-4^. However, the molecular determinants of 2C-like conversion have not been completely elucidated. Here, we show that CTCF is a barrier for 2C-like reprogramming. Indeed, forced conversion to a 2C-like state by DUX expression was associated with DNA damage at a subset of CTCF binding sites. Endogenous or DUX-induced 2C-like ESC showed decreased CTCF enrichment at known binding sites, suggesting that acquisition of a totipotent-like state is associated with a highly dynamic chromatin architecture. Accordingly, depletion of CTCF in ESC efficiently promoted spontaneous and asynchronous conversion to a totipotent-like state. This phenotypic reprogramming was reversible upon restoration of CTCF levels. Furthermore, we showed that transcriptional activation of the ZSCAN4 cluster was necessary for successful 2C-like reprogramming. In summary, we revealed the intimate relation between CTCF and totipotent-like reprogramming.

---

Totipotency is defined as the ability of a single cell to generate all cell types and is found in zygotes and 2-cell (2C) embryos^1,2^. As development proceeds, embryonic cells progressively restrict their developmental potential. Embryonic stem cells (ESC) isolated from the inner cell mass (ICM) of blastocysts are defined as pluripotent since they lack the ability to differentiate into extra-embryonic tissues^1,2^. Interestingly, a rare (∼1-2%) transient population of cells with totipotent-like potential was identified within ESC cultures^2-4^. This cell population expresses high levels of transcripts detected in 2C embryos, including a specific gene set regulated by endogenous retroviral promoters of the MERVL subfamily^2-4^. At the 2C embryonic stage, these retroviral genetic elements are re-activated and highly expressed when the zygotic genome is first transcribed and quickly silenced after further development. Based on this specific feature, retroviral promoter sequences (*LTR*) have been used as a reporter system to genetically label 2C-like cells *in vitro* to study their behavior and properties^2-4^. Previous studies have shown the role of different genes and pathways in converting ESC to a totipotent-like state *in vitro*^3,4^. Indeed, expression of the transcription factor DUX in ESC is necessary and sufficient to induce a 2C-like conversion characterized by similar transcriptional and chromatin accessibility profiles, including MERVL activation, as observed in 2C-blastomeres^5-7^. This reprogramming cell model has been instrumental to study the molecular mechanisms that regulate the acquisition and maintenance of totipotent-like features. DUX belongs to the double homeobox family of transcription factors exclusive to placental mammals^8^ and is expressed exclusively in the 2C embryo^5-7^. Interestingly, DUX knockout mice revealed that DUX is important but not essential for development, suggesting that additional mechanisms regulate zygotic genome activation (ZGA) and the associated totipotent state *in vivo*^9,10^.

## 2C-like conversion correlates with DNA damage and cell death

To explore new molecular determinants regulating totipotency, we generated ESC carrying a doxycycline (DOX)-inducible DUX cDNA (hereafter, ESC^Dux^)^11^. Upon DOX activation we detected the expected expression of *Dux* and its downstream ZGA-associated genes (Extended data Fig. 1a, b). In addition, ESC^Dux^ containing an *LTR-RFP* reporter showed reactivation of MERVL sequences after DOX induction (Extended data Fig. 1c, d). Over-expression of DUX triggers toxicity in myoblasts^12^. However, whether sustained expression of DUX leads to cell death in ESC has not been explored thoroughly. We observed that DUX expression induced cell death in a dose and time-dependent manner and correlated with the extent of 2C-like conversion (Fig. 1a). Indeed, live cell imaging of DOX-treated ESC^Dux^ expressing H2B-eGFP showed efficient cell death in cells asynchronously converting to a 2C-like state (Fig. 1b, Supplementary Video 1). Interestingly, accumulation of DOX-induced ESC^Dux^ in the G1 and G2 phases of the cell cycle along with a decrease in DNA replication preceded cell death (Fig. 1c, d). To exclude that these effects were due to supra-physiological levels of DUX, we analyzed the unperturbed subpopulation of ESC that spontaneously undergoes a 2C-like conversion^**3**^. These endogenous totipotent-like ESC were also characterized by G2 accumulation, decreased DNA replication, and overt spontaneous cell death following 2C-like conversion (Extended data Fig. 2a-c and Supplementary Video 2). In support of these observations, the activation of the transcriptional 2C program during ZGA following the first cleavage in fertilized zygotes is accompanied by an extremely long G2 phase (around 12-16 hours)^13,14^.

**Fig. 1:**
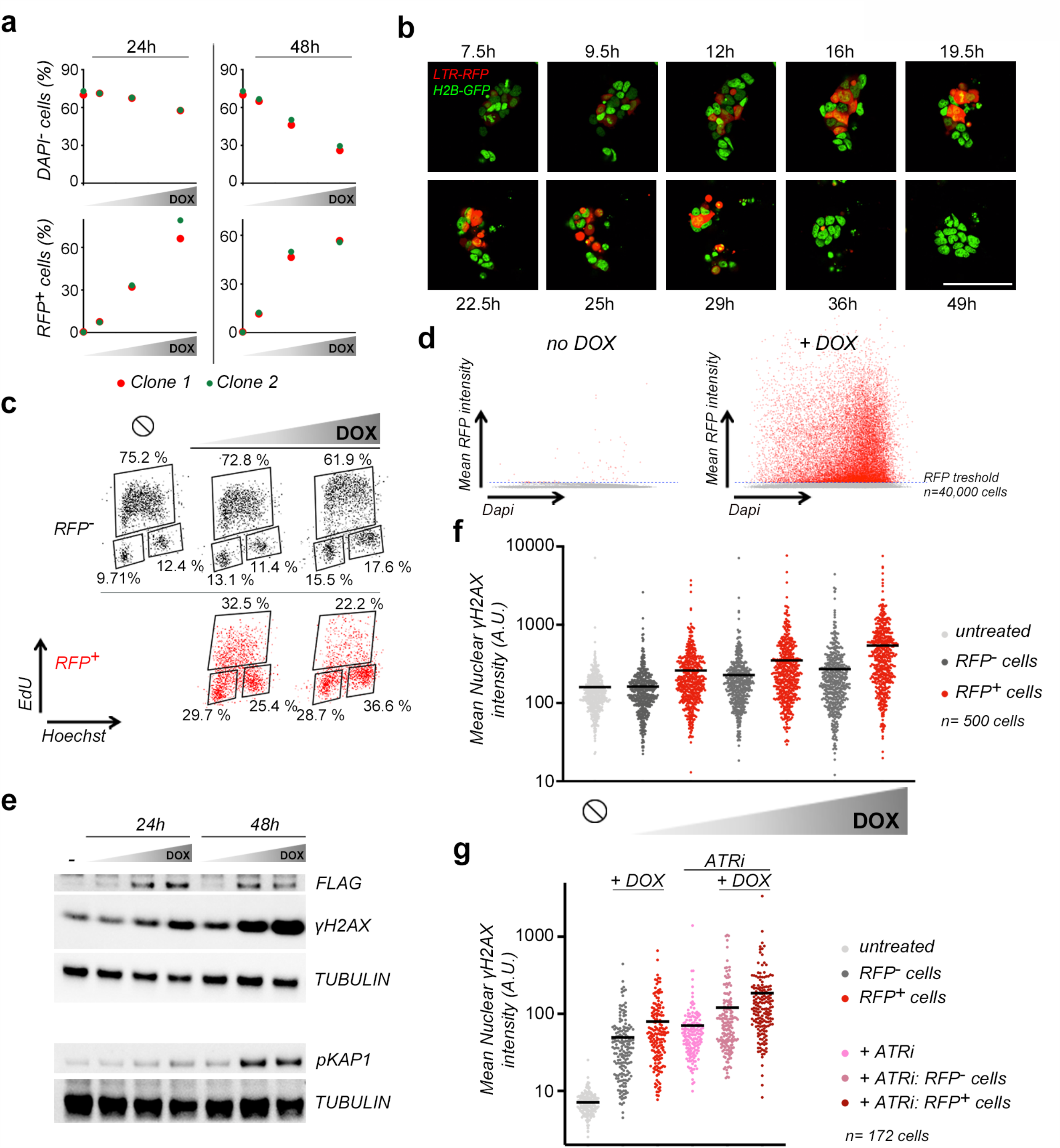
Induction of totipotent-like features in ESC correlated with DNA damage and cell death. **a**, Plots showing the percentage of alive (DAPI^-^) and RFP^+^ cells from *LTR-RFP* reporter ESC^DUX^ treated with increasing doses of DOX (150, 300 and 600 ng/ml) for the indicated time points. Data was collected by flow cytometry. **b**, Representative images obtained from a time lapse experiment where *LTR-RFP* reporter ESC^DUX^ expressing H2B-eGFP were treated with DOX and imaged at the indicated timepoints. Scale bar, 100 µm. **c**, Flow cytometry analysis of the cell cycle distribution in untreated or DOX-treated *LTR-RFP* reporter ESC^DUX^ for 24 hours. Percentages for each phase of the cell cycle are included. **d**, Dot plots showing cell distribution based on DNA content and RFP expression in *LTR-RFP* reporter ESC^DUX^ treated with DOX for 24 hours. **e**, Western blot analysis of the indicated proteins performed in ESC^DUX^ treated with different doses of DOX for the indicated time points. Expression of DUX was monitored by its FLAG-tag. Tubulin levels are shown as a loading control. **f**, High-throughput imaging (HTI) quantification of γH2AX in *LTR-RFP* reporter ESC^DUX^ treated with different concentrations of DOX for 24 hours. Center lines indicate mean values. ∅=No treatment. **g**, HTI quantification of γH2AX in *LTR-RFP* reporter ESC^DUX^ treated with DOX and/or with 1µM ATR inhibitor (ATRi) for 24 hours. Center lines indicate mean values. In (**c**), (**f**) and (**g**), cells were split in RFP^-^ or RFP^+^. In (**a-g**) data are representative of at least two independent experiments performed in two different clones.

We next examined whether G2 accumulation correlated with DUX-induced DNA damage. We observed that sustained expression of DUX leads to DNA-damage, revealed by the increased levels of phosphorylated H2AX (γH2AX) and KRAB-associated protein 1 (KAP1) in a dose and time-dependent manner (Fig. 1e, f). We also detected higher levels of γH2AX in endogenous 2C-like ESC (Extended data Fig. 2d). The decrease in DNA replication and elevated levels of γH2AX observed in 2C-like ESC suggested that replication stress (RS) could underlie the increased levels of DNA damage in these cells. Accordingly, increasing RS levels by using an ATR inhibitor showed an additive effect of RS and DUX expression on DNA damage (Fig. 1g). Our results showed that induction of a totipotent-like state in ESC induced G2 accumulation and decreased cell viability associated with replication stress-mediated DNA-damage.

## Reduced levels of chromatin bound CTCF in 2C-like ESC

We next sought to investigate the nature of the DNA damage. Since DUX is a potent transcriptional activator, we hypothesized that RS-induced DNA damage was localized in specific regions of the genome rather than being randomly distributed. To explore this possibility, we performed END-seq^15^, a highly sensitive method to detect DNA ends (single or double strand breaks) genome-wide at base-pair resolution. DUX-expressing ESC showed increased accumulation of ENDseq signal compared to untreated ESC^Dux^ (Fig. 2a). A total of 1539 ENDseq peaks overlapped between two independent ESC^Dux^ clones (Supplementary Tables 1-3). Moreover, the type of lesion (double or single strand DNA break) at each site, showed high correlation when both ESC^Dux^ clones were compared (Extended data Fig. 3a-c). More than 25% of the ENDseq peaks localized within a 10kb distance from a DUX binding site. Furthermore, 16% of the 1220 genes associated by proximity to ENDseq peaks, including well-known 2C genes, were strongly upregulated by DUX (Extended data Fig. 3d and Supplementary Tables 4, 5). These results showed that DUX-induced 2C-like conversion reproducibly generated DNA lesions in specific genomic regions associated with DUX-induced transcription. We next asked whether these regions shared any feature that could explain the reiterative DNA damage on them. Thus, we performed a transcription factor motif enrichment analysis using our END-seq peak dataset and found the CTCF binding motif as one of the most significant (Extended data Fig. 3e). Using published CTCF ChIPseq datasets in ESC^16^, we confirmed that around 50% of the END-seq peaks were occupied by CTCF (Fig. 2b, c, Extended data Fig. 3c, f, g and Supplementary table 6). Moreover, these sites were also enriched in SMC1 and SMC3^17^, components of the cohesin ring-like protein complex (Fig. 2b, c, Extended data Fig. 3f).

**Fig. 2:**
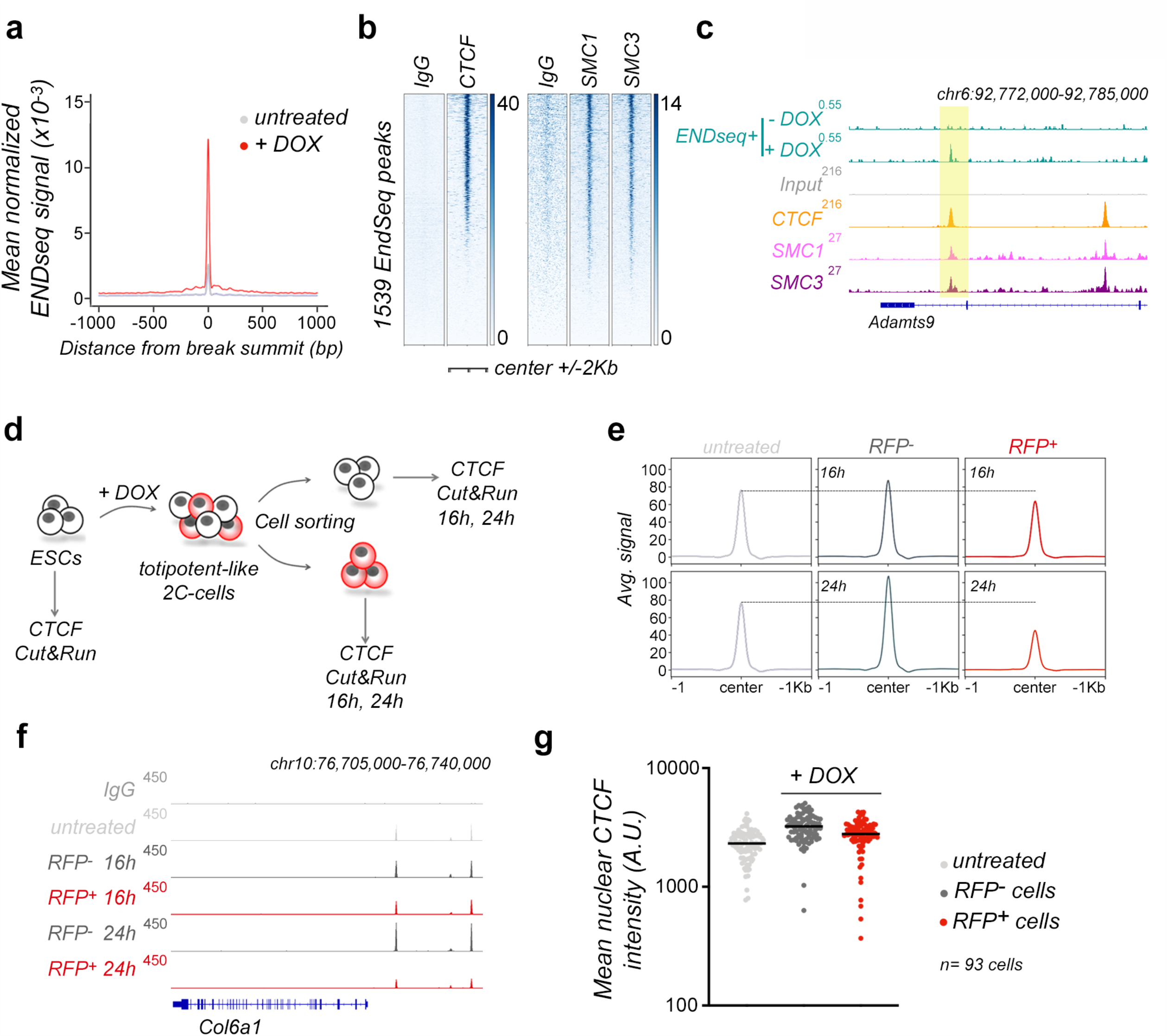
2C-like ESC are characterized by decreased levels of chromatin-bound CTCF. **a**, Plot showing aggregated ENDseq signal in untreated and DOX-treated ESC^DUX^ for 16 hours from one clonal line in the set of overlapped 1539 ENDseq sites identified in two independent DOX-treated ESC^DUX^ clones. **b**, Heatmaps showing CTCF^20^, SMC1 and SMC3^21^ occupancy at the set of 1539 ENDseq sites. **c**, Genome browser tracks showing ENDseq signal in untreated and DOX-treated ESC^DUX^ at the indicated genome location. In addition, CTCF^20^, SMC1 and SMC3^21^ occupancy in ESC is shown. ENDseq peak is highlighted. **d**, Schematic representation of the experiment performed. **e**, Cut&Run read density plot (RPKM) showing CTCF occupancy in the set of 50183 CTCF sites identified in ESC^20^ in the cell samples shown in (**d**). The signal obtained in corresponding inputs (IgG) was subtracted. **f**, Genome browser tracks showing CTCF occupancy at the indicated genome location in the cell samples shown in (**d**). **g**, HTI quantification of CTCF in untreated or DOX-treated ESC^DUX^ for 24 hours. Cells were split into RFP^+^ or RFP^-^ subpopulations. Center lines indicate mean values. In (**a**) and (**e-g**) representative data from one ESC^DUX^ clone is shown but two independent clones were analyzed. In (**b, c** and **f**) input (IgG) is shown as a background reference control.

The transcription factor CTCF is a zinc-finger binding protein involved in chromosome folding and insulation of topologically associated domains (TADs)^18^. Based on the observed CTCF-associated DNA damage in DOX-induced ESC^Dux^, we speculated that CTCF might represent a barrier for the reprogramming to totipotency. This idea was supported by two observations. First, cohesin depletion in differentiated cells facilitates reprogramming during somatic cell nuclear transfer by activating ZGA^19^. Second, totipotent zygotes and 2C embryos are characterized by chromatin in a relaxed state associated with weak TADs^20-21^. Following fertilization, development is accompanied by a progressive maturation of high-order chromatin architecture^20-21^. Interestingly, increasing levels of CTCF during human ED are required for the progressive establishment of TADs^22^. Similarly, we also observed a steady increase in the levels of CTCF during development in mouse embryos (Extended data Fig. 4). To examine whether levels of chromatin-bound CTCF correlated with totipotency features, we first assessed the CTCF binding landscape in 2C-like cells by native Cut&Run sequencing. For this, we used *LTR-RFP* reporter ESC^DUX^ to first induce 2C-like conversion, and then, sort RFP^+^ and RFP^-^ cells at two different timepoints, 16 and 24 hours after DOX induction (Fig. 2d-f). Interestingly, 16 hours after DOX induction, RFP^-^ ESC showed a slight increased CTCF enrichment at known CTCF sites^20^ compared to non-induced ESC while RFP^+^ showed the opposite trend (Fig. 2e, f). Changes in CTCF enrichment were further enhanced in cells reprogrammed 24 hours after DUX expression (Fig. 2e, f) and were not due to variations in the total levels of CTCF (Fig. 2g). Significantly, spontaneously converting 2C-like ESC showed a similar reduction in CTCF enrichment at known sites^20^ compared to pluripotent ESC (Extended data Fig. 5). Combined, these results demonstrated that totipotent-like cells are characterized by decreased levels of chromatin-bound CTCF, indicative of a more relaxed chromatin architecture.

## CTCF depletion leads to spontaneous 2C-like conversion

To examine whether reduced levels of CTCF enrichment are causative for acquiring totipotency-like features, we used an auxin-inducible degron system to deplete CTCF in ESC^23^. This cell line (ESC^CTCF-AID^ hereafter) harbors both *Ctcf* alleles tagged with an auxin-inducible degron (AID)^24^ sequence fused to eGFP. Although CTCF-AID protein levels in ESC^CTCF-AID^ are lower compared to untagged CTCF in wild-type cells, ESC^CTCF-AID^ showed negligible transcriptional changes as tagged CTCF retains most functionality^23^. To test whether CTCF deletion induces conversion to 2C-like cells we first examined in CTCF-depleted cells the expression levels of the zinc finger protein ZSCAN4, a gene cluster that is selectively expressed in 2C embryos and 2C-like ESC^3,25^. Strikingly, ZSCAN4 levels were elevated two days following CTCF depletion and further increased two days later (Fig. 3a and Extended data Fig. 6a, b). Indeed, more than 20% of the cells expressed ZSCAN4 three days following CTCF depletion (Extended data Fig. 6b). Importantly, similar percentages of RFP^+^ cells were observed in *LTR-RFP* reporter ESC^CTCF-AID^ (Fig. 3b). This percentage decreased upon restoration of the CTCF levels by washing off auxin (Fig. 3b). Using RNAseq datasets from CTCF-depleted cells at different timepoints, we observed a progressive increase in the expression of genes enriched or exclusively expressed in 2C embryos or 2C-like ESC (Fig. 3c, d and Supplementary Table 7). Among these, endogenous MERVL sequences as well as *Dux* were selectively expressed over time upon CTCF depletion (Fig. 3d). We also observed decreased expression of the pluripotent gene OCT4 in ZSCAN4^+^ auxin-treated ESC^CTCF-AID^ as described for 2C-like ESC (Fig. 3e)^5^. Furthermore, CTCF-depleted ESC showed transcriptional similarity with DUX-overexpressing ESC (Extended data Fig. 6c). In addition, 2C-like reprogramming was further boosted cooperatively by expressing low levels of DUX or by incubating ESC^CTCF-AID^ with HDAC inhibitors, known to promote 2C-like conversion (Extended data Fig. 6d, e). Finally, we validated these observations by generating additional ESC^CTCF-AID^ clonal lines (Extended data Fig. 6f). Collectively, these results demonstrated that CTCF depletion leads to spontaneous 2C-like conversion in ESC.

**Figure 3:**
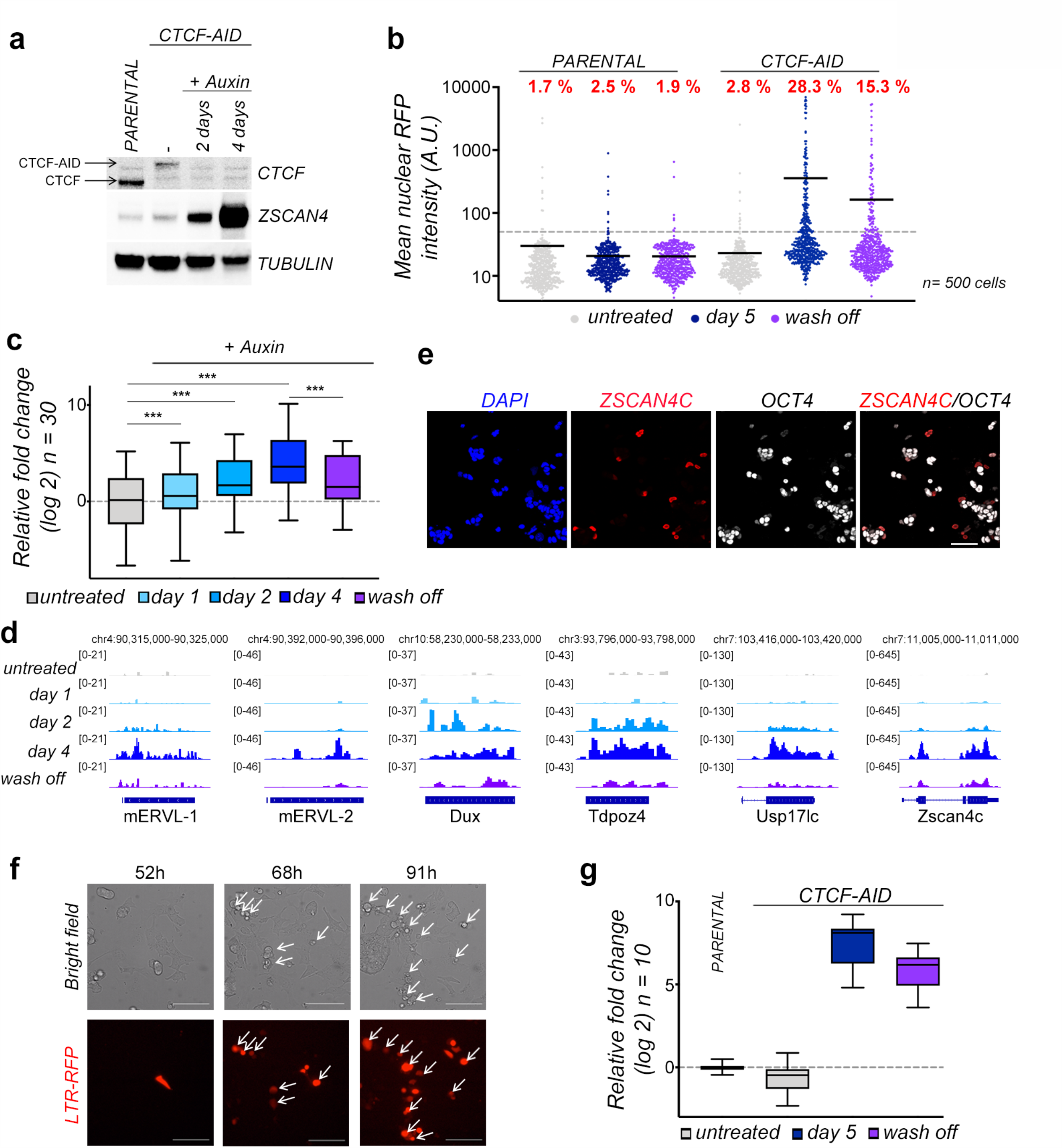
Spontaneous 2C-like conversion in CTCF-depleted ESC. **a**, Western blot analysis of the indicated proteins performed in ESC^CTCF-AID^ treated with auxin for two and four days. Parental ESC were used to show the smaller size and higher levels of CTCF compared to ESC^CTCF-AID^. Tubulin levels are shown as a loading control. **b**, HTI quantification of RFP^+^ cells in untreated or auxin-treated for five days *LTR-RFP* reporter parental ESC and ESC^CTCF-AID^. RFP^+^ cells two days after a wash off following three days of auxin treatment were also quantified. Center lines indicate mean values. Percentages of RFP^+^ cells above the threshold are indicated. **c**, Graph showing the relative fold change (log2) expression of a subset of thirty 2C associated genes in ESC^CTCF-AID^ treated with auxin for one, two and four days (Supplementary Table 7). Untreated and wash off ESC^CTCF-AID^ were also included. Data was obtained from RNAseq datasets^23^. **d**, Genome browser tracks showing RNAseq RPKM read count at the indicated genes in the same samples as in (**c**). **e)** Immunofluorescence analysis of ZSCAN4 and OCT4 in ESC^CTCF-AID^ treated with auxin for 4 days. DAPI was used to visualize nuclei. Scale bars, 100 µm. **f**, Representative bright field images (upper panels) obtained from a time lapse experiment performed in ESC^CTCF-AID^ treated with auxin. RFP^+^ cells are shown as they convert over time (lower panels). Time since the addition of auxin is indicated. White arrows indicate 2C converted cells undergoing cell death. Scale bars, 100 µm. **g**, Graph showing the relative fold change (log2) expression of a subset of ten 2C associated genes (DUX, ZSCAN4, ZFP352, TCSTV3, SP110, TDPOZ1, DUB1, EF1a, PRAMEL7 and MERVLs) in *LTR-RFP* reporter ESC^CTCF-AID^ untreated or treated with auxin for three days and further incubated with auxin or washed off for additional 18 hours (a total of five days) and sorted based on RFP expression. Parental ESC were also sorted and included as a reference. GAPDH expression was used to normalize gene expression. In (**a, b** and **f**), one representative experiment is shown but at least two independent experiments were performed.

We next examined the dynamics of the 2C-like conversion by live cell imaging in *LTR-RFP* reporter ESC^CTCF-AID^. Reprogramming to 2C-like ESC is asynchronous as ESC convert over time after CTCF depletion (Fig. 3f). Interestingly, we observed that spontaneously converted 2C-like ESC undergo similar cell death as shown for endogenous 2C-like ESC while non-converted ESC divide and do not show overt cell death (Fig. 3f, Supplementary Video 3). Accordingly, CTCF-depleted 2C-like ESC showed increased γH2AX, similar to endogenous 2C-like ESC (Extended data Fig. 6g). Our data suggested that cell toxicity induced by CTCF-depletion is due to the selective death of the spontaneously converted 2C-like ESC. Finally, we explored whether restoring CTCF expression facilitates the exit from the totipotent-like state. For this, CTCF-depleted *LTR-RFP* reporter ESC^CTCF-AID^ for four days were either further incubated with auxin or washed off for an additional 18 hours (5 days total) and sorted based on RFP expression. Gene expression analysis showed that restoration of CTCF levels induced a decrease in the 2C-like transcriptional program in 2C-like cells anticipating the exit from the totipotent-like state (Fig. 3g and Extended data Fig. 7). Collectively, these results demonstrated that chromatin bound CTCF prevents 2C-like conversion.

## ZSCAN4 expression is required for 2C-like reprogramming

Endogenous emergence of 2C-like cells in ESC cultures is a stepwise process defined by sequential changes in gene expression^26^. ZSCAN4^+^MERVL^-^ ESC are detected during this process and represent an intermediate step that precedes the full conversion to a 2C-like state^26,27^. Levels of ZSCAN4 progressively increase during 2C conversion prior to the activation of MERVL sequences and the expression of chimeric transcripts^26,27^. We also detected a progressive accumulation of ZSCAN4 in CTCF-depleted ESC starting as early as 24 hours after depletion (Fig. 4a and Extended data Fig. 8a, b). However, upregulation of DUX or MERVL sequences was observed at later timepoints, suggesting that spontaneous conversion upon CTCF depletion followed a similar molecular roadmap as endogenous 2C-like cells. In agreement, we also detected ZSCAN4^+^mERVL^-^ ESC in early auxin treated *LTR-RFP* reporter ESC^CTCF-AID^ (Extended data Fig. 8c). We next asked whether early transcriptional activation of ZSCAN4 in ESC precursors is essential for full conversion to 2C-like cells. Therefore, we infected *LTR-RFP* reporter ESC^CTCF-AID^ with lentiviruses expressing shRNAs against ZSCAN4 and examined transcriptional dynamics and 2C-like conversion upon CTCF removal^28^. Surprisingly, downregulation of ZSCAN4 in CTCF-depleted cells impaired expression of 2C markers and abrogated reprogramming to 2C-like cells (Fig. 4b, c and Extended data Fig. 8d). Furthermore, over-expression of ZSCAN4C boosted 2C-like conversion as early as 24 hours specifically in CTCF-depleted ESC while cells with normal levels of CTCF did not show major changes in the number of 2C-like cells (Fig. 4d and Extended data Fig. 8e). These combined results demonstrated that ZSCAN4 proteins are essential for the 2C-like conversion mediated by CTCF depletion.

**Figure 4:**
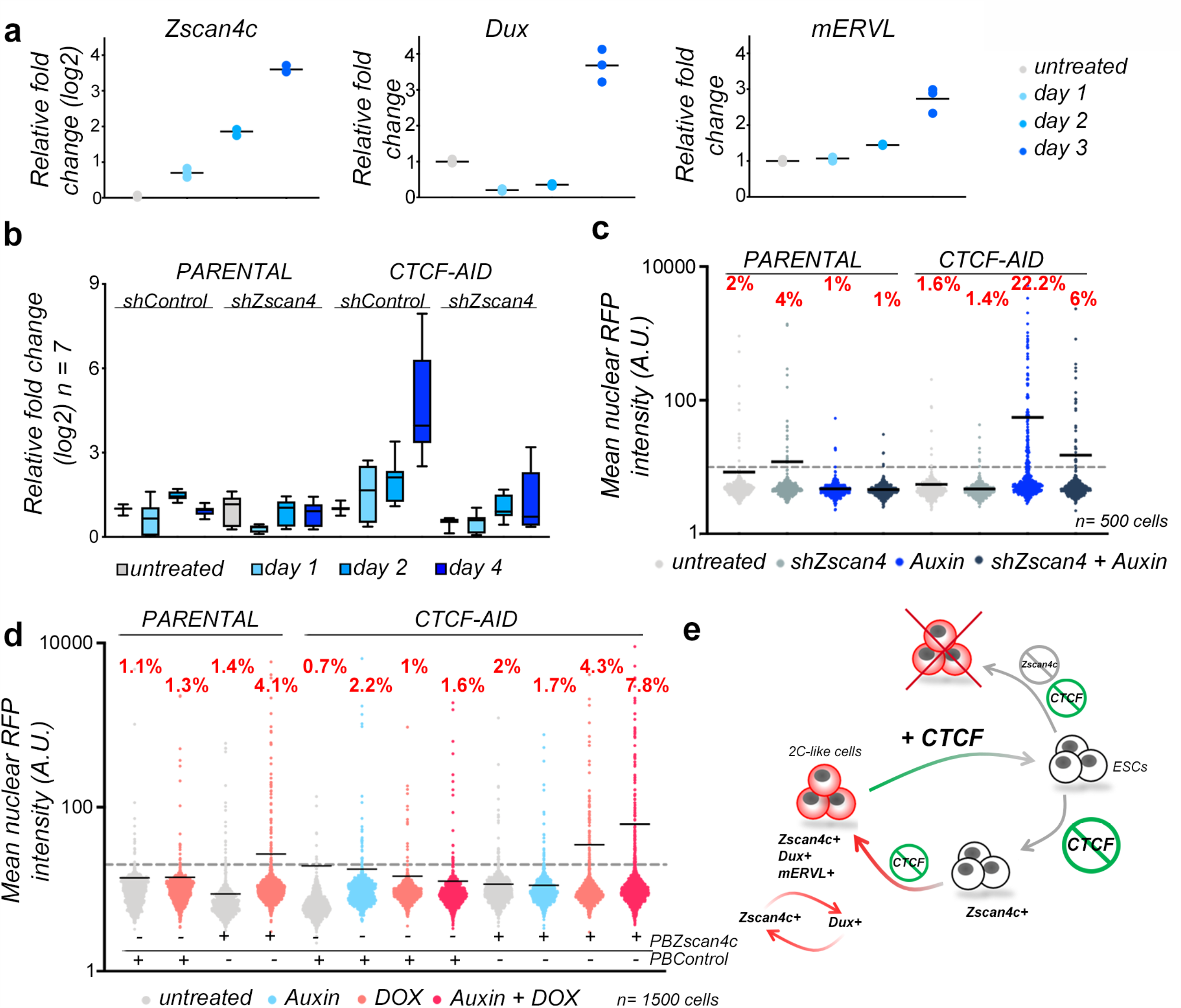
Transcriptional activation of the ZSCAN4 cluster is required for 2C-like reprogramming. **a**, Graph showing the relative fold change (log2 or linear) expression of DUX, ZSCAN4 and MERVL in untreated or auxin-treated *LTR-RFP* reporter ESC^CTCF-AID^ for the indicated days. Data are shown by triplicate. **b**, Graph showing the averaged relative fold change (log2) expression of seven 2C genes (DUX, ZSCAN4, ZFP352, TCSTV3, SP110, TDPOZ1 and MERVLs) in untreated or auxin-treated at the indicated time points in *LTR-RFP* reporter control and ESC^CTCF-AID^. ESC were infected with lentiviruses expressing control or shRNAs against ZSCAN4. Reactions were performed by triplicate in two independent experiments. **c**, HTI quantification of RFP^+^ cells in untreated or auxin-treated for four days *LTR-RFP* reporter control and ESC^CTCF-AID^. ESC were infected with lentiviruses expressing control or shRNAs against ZSCAN4. Center lines indicate mean values. **d**, HTI quantification of RFP^+^ cells in untreated or auxin-treated for 24 hours *LTR-RFP* reporter control and ESC^CTCF-AID^. ESC harbor a DOX-inducible piggyBac (PB) construct expressing ZSCAN4C and were induced as indicated together with auxin. Center lines indicate mean values. **e**, Schematic representation of the model inferred from the data presented here. In (**a-d**), one representative experiment is shown but at least two independent experiments were performed. In (**c, d**) percentages of RFP^+^ cells above the threshold are indicated.

## DISCUSSION

Our study demonstrates that 2C-like ESC are unstable *in vitro*. We observed increased DNA damage and cell death in endogenous, DUX-induced and CTCF-depleted 2C-like ESC. Similarly, over-expression of DUX *in vivo* leads to developmental arrest and embryo death ^29^. We show that the DNA damage observed in DUX-induced 2C-like ESC might be at least partially associated to replication stress and involves the generation of single or double strand brakes at certain CTCF sites. We speculate that proximity of ENDseq peaks to DUX-binding sites might induce local *de novo* transcription/replication conflicts. Inefficient release of nearby bound CTCF in ESC undergoing 2C-like conversion could promote fork stalling and eventual breakage. Further work will be needed to understand the exact origin of these DNA breaks. Nevertheless, additional sources of damage are likely to be associated with the 2C-like state or induced by DUX. In fact, human ortholog DUX4 mediates the accumulation of dsRNA foci and the activation of the dsRNA response contributing to the apoptotic phenotype associated with DUX over-expression^30^.

CTCF depletion triggers spontaneous 2C-like conversion and promotes the acquisition of totipotent-like features in ESC (Fig. 4e). Importantly, expression of the ZSCAN4 gene cluster is a necessary early event in this conversion and, although its precise role in this process is unclear, ZSCAN4 has been implicated in protecting the 2C embryo from DNA damage^28,31^. Thus, ZSCAN4 could participate in limiting the damage associated with the 2C-like conversion. Expression of DUX, which is a later event, enhances the transcriptional activation of the ZSCAN4 cluster by direct DUX binding to its promoters. In fact, DUX knockout ESC and embryos showed defective ZSCAN4 activation. This positive feedback loop might be required to stabilize the 2C-like state^9-10^.

Totipotent cells display high core histone mobility compared to pluripotent cells^32, 33^. Similarly, our results indicate that totipotency is associated with dynamic chromatin architecture characterized by decreased levels of chromatin-bound CTCF. CTCF binds to a large number of endogenous RNAs and this interaction seems important for chromatin CTCF deposition^34^. Indeed, CTCF mutants unable to bind RNA showed decreased genome-wide binding^34^. It is tempting to speculate that the progressive strength of TADs during ED^20-21^ correlates with increasing levels of CTCF and RNA transcription after ZGA. Further work will be needed to address how CTCF deposition and TAD insulation take place during early development and if these events play an active role in promoting the exit from totipotency in the early embryo. In summary, we revealed the intertwined relation between CTCF and totipotent-associated features.

## METHODS

### Embryo Culture

C57BL/6J mice were obtained from the Jackson Laboratory. All the animal work included here was performed in compliance with the NIH Animal Care & Use Committee (ACUC) Guideline for Breeding and Weaning. For embryo isolation, 4-weeks old female mice were injected intraperitoneally with 5IU Pregnant Mare Serum Gonadotropin (PMSG, Prospec) followed by 5 IU human Chorionic Gonadotropin (hCG, Sigma-Aldrich) 46-48 hours later. Pregnant females were euthanized, and embryos collected in M2 media (MR-015-D, Sigma-Aldrich) at indicated time points after hCG injection: E0.5, E1.0, E2.5 and E3.5. The sex of embryos was not determined. Isolated embryos were fixed for 10 min in 4% Paraformaldehyde (Electron Microscopy Sciences), permeabilized for 30 min in 0.3% Triton X-100 and 0.1M Glycine in PBS 1X and blocked for 1 hour (1% BSA, 0.1% Tween in PBS 1X), followed by overnight incubation with primary antibodies against CTCF (1:1000 dilution, ab188408, Abcam). Embryos were washed in 0.1% Tween in PBS 1X and incubated with the appropriate secondary antibody for 1 hour at room temperature. Embryos were imaged using a Nikon Ti2-E microscope (Nikon Instruments) equipped with a Yokogawa CSU-W1 spinning diskunit, a Photometrics BSI sCMOS camera and 20x (N.A. 0.75) and 60x (N.A. 1.49) plan-apochromat objective lenses. Confocal z-stacks were acquired and used to generate 3D surfaces were rendered based on nuclear DAPI-staining and the corresponding regions were used to quantify the fluorescence intensity of CTCF. Embryo z-stack images were quantified using Imaris Bitplane (Oxford Instruments).

### Cell culture

Wild-type (R1 and G4) ESC, ESC^DUX^ and ESC^CTCF-AID^ (ID: EN52.9.1)^23^ were grown on a feeder layer of growth-arrested MEFs or on gelatin 0.1% in high-glucose DMEM (Invitrogen) supplemented with 15% FBS, 1:500 LIF (made in house), 0.1 mM nonessential amino acids, 1% glutamax, 1mM Sodium Pyruvate, 55 mM β-mercaptoethanol, and 1% penicillin/streptomycin (all from Life Technologies) at 37°C and 5% CO_2_. Cells were routinely passaged with Trypsin 0.05% (Gibco). Media was changed every other day and passaged every 2-3 days. HEK293T (American Type Culture Collection) cells were grown in DMEM, 10% FBS, and 1% penicillin/streptomycin. Generation of infective lentiviral particles and ESC infections were performed as described^35^.

To generate ESC^DUX^ cell lines, a FLAG-tag version of the codon-optimized mouse DUX was amplified by PCR (Primers in Extended Table 1) from pCW57.1-mDUX-CA (Addgene 99284) and subcloned into the pBS31 plasmid (pBS31-FLAG_mDUX). A Flp-dependent recombination event using pBS31-FLAG_mDUX in the KH2 ESC line was used to knock-in the cDNA for FLAG_mDUX into a tetO-minimal promoter allocated in the *Col1a1* locus as described^11^.

To generate additional ESC^CTCF-AID^ cell lines, R1 and ESC^DUX^ were co-transfected using jetPRIME (PolyPlus transfection) with the plasmids CTCF-AID[71-114]-eGFP-FRT-Blast-FRT (92140, Addgene), pCAGGS-Tir1-V5-BpA-Frt-PGK-EM7-NeoR-bpA-Frt-Rosa26 (92140, Addgene) and the plasmid pX330-U6-Chimeric_BB-CBh-hSpCas9 (42330, Addgene) encoding sgRNAs targeting CTCF and ROSA26 alleles (see Extended Table 1 for sgRNA sequences). Two days after transfection ESC were selected with Neomycin (200ug/ml) for one additional week. Individual ESC clones were picked and amplified based on eGFP expression indicating successful CTCF targeting. HTI and western blot analyses were used to verify that eGFP and CTCF were lost upon addition of 500 µM auxin for 24 hours.

To generate ESC lines carrying the *LTR-RFP* reporter, the LTR sequence was PCR amplified and subcloned in a piggyBac plasmid upstream of a turboRFP (RFP) coding region to generate the *LTR-RFP* reporter (Primers in Extended Table 1). PiggyBac-*LTR-RFP* plasmid together with a plasmid encoding for a supertransposase were co-transfected in ESC and further selected with Neomycin (200ug/ml) for one week. To generate ESC^CTCF-AID^ lines carrying a DOX-inducible ZSCAN4-PiggyBac construct, the coding sequence for ZSCAN4C was amplified from cDNA and subcloned into the plasmid PB-TRE-dCas9-VPR (63800, Addgene), after removing the dCas9-VPR insert. DOX-inducible PiggyBac-*ZSCAN4C* plasmid together with a plasmid encoding for a supertransposase were co-transfected in ESC and further selected with Hygromycin (200ug/ml) for one week. To generate ZSCAN4-knockdown ESC^CTCF-AID^ lines, cells were infected with pLKO.1 control or pLKO.1-shZSCAN4 (5’-GAATGCAACAACTCTTGTAATCTCGAGATTACAAGAGTTGTTGCATTCT-3’, Millipore Sigma) and further selected with Puromycin (1ug/ml) for one week.

### Immunofluorescence

Cells were fixed in 4 % Paraformaldehyde (PFA, Electron Microscopy Sciences) for 10 min at RT followed by 10 min of permeabilization using the following permeabilization buffer (100 mM Tris-HCl pH 7.4, 50 mM EDTA pH 8.0, 0.5 % Triton X-100). The following primary antibodies were incubated overnight: OCT3/4 (1:100, sc-5279, Santa Cruz Biotechnology), ZSCAN4 (1:2000, AB4340, Millipore Sigma), γH2AX (1:1000, 05-636, Millipore), CTCF (1:1000, ab188408, Abcam), Flag (1:500, F1804, Sigma Aldrich). Corresponding Alexa-Fluor (−488, −568 and −647) secondary antibodies were used to reveal primary antibody binding (Thermo Fisher Scientific). For generating the plots shown in Figure 1d, image analysis was performed using a custom Python script. In brief, DAPI-stained nuclei were segmented using the StarDist deep-learning image segmentation^36^. Segmented nuclei ROIs were used to quantify total DAPI intensity and RFP mean intensity.

### High throughput imaging (HTI)

A total of 10,000-20,000 ESC (depending on the experiment and on the specific ESC line) were plated on gelatinized μCLEAR bottom 96-well plates (Greiner Bio-One, 655087). ESC were treated with DOX (different concentrations in the range from 150–600 ng/ml) or 500 µM auxin as indicated or incubate with 10µM EdU (Click Chemistry Tools) for 30 minutes before fixation with 4% PFA in PBS for 10 minutes at room temperature. γH2AX and ZSCAN4 staining was performed using standard procedures. EdU incorporation was visualized using Alexa Fluor 488-azide or Alexa Fluor 647-azide (Click Chemistry Tools) Click-iT labeling chemistry and DNA was stained using DAPI (4’,6-diamidino-2-phenylindole). When indicated, ESC^DUX^ were treated with 1 µM ATR inhibitor (AZ20, Selleckchem).

Cooperation between CTCF-depletion and DUX expression was examined in CTCF-AID targeted ESC^DUX^ upon treatment with auxin and low concentration of DOX. Similarly, Cooperation between CTCF-depletion and HDAC inhibition was examined in ESC^CTCF-AID^ treated with auxin and 10 µM HDAC inhibitor.

Images were automatically acquired using a CellVoyager CV7000 high throughput spinning disk confocal microscope (Yokogawa, Japan). Each condition was performed in triplicate wells and at least 9 different fields of view (FOV) were acquired per well. High-Content Image (HCI) analysis was performed using the Columbus software (PerkinElmer). In brief, nuclei were first segmented using the DAPI channel. Mean fluorescence intensities for γH2AX, ZSCAN4, CTCF, eGFP or RFP signal were calculated over the nuclear masks in their respective channels. Single cell data obtained from the Columbus software was exported as flat tabular .txt files, and then analyzed using RStudio version 1.2.5001, and plotted using Graphpad Prism version 9.0.0.

### Live Cell imaging

When indicated, ESC were infected with a lentiviral plasmid encoding H2B-GFP (kind gift from Marcos Malumbres, CNIO, Spain). A total of 40,000 ESC were plated in gelatine-coated µ-Slide 8 wells plates (80826, Ibidi) and imaged untreated or Auxin/DOX-treated for a time period between 43-48hrs depending on the experiment. Images were acquired every 15 or 20 minutes over the time course using either a Nikon spinning disk confocal microscope or a Zeiss LSM780 confocal microscope equipped with 20x plan-apochromat objective lenses (N.A. 0.75 and 0.8, respectively) and stage top incubators to maintain temperature, humidity and CO2 (Tokai Hit STX and Okolab Bold Line, respectively).

### Western blot

Trypsinized cells were lysed in 50 mM Tris pH 8, 8 M Urea (Sigma) and 1% Chaps (Millipore) followed by 30 min of shaking at 4°C. 20 μg of supernatants were run on 4%-12% NuPage Bis-Tris Gel (Invitrogen) and transferred onto Nitrocellulose Blotting Membrane (GE Healthcare). Membranes were incubated with the following primary antibodies overnight at 4°C: p-KAP1 (dilution 1:1000, A300-767A, Bethyl) or ZSCAN4C (1:500, AB4340, Millipore Sigma), γH2AX (1:1000, 05-636, Millipore), CTCF (1:1000, 07-729, Millipore), Flag (1:1000, F1804, Sigma Aldrich), Tubulin (1:50000, T9026, Sigma-Aldrich). The next day the membranes were incubated with HRP-conjugated secondary antibodies (1:5000) for 1 h at room temperature. Membranes were developed using SuperSignal West Pico PLUS (Thermo Scientific).

### Flow cytometry and cell sorting

For live cell flow cytometry experiments, cells were dissociated into single cell suspensions and analyzed for RFP expression, DAPI was added to detect cells with compromised membrane integrity. For EdU Click-IT experiments, cells were incubated for 20 min with 10 µM EdU, fixed in 4 % paraformaldehyde, permeabilized in 0.5 % triton X-100, followed by Alexa Flour 488-azide or Alexa Flour 647-azide Click-iT labeling chemistry. DNA content was stained using DAPI or Hoechst 33342 (62249, Thermo Fisher Scientific). Analytic flow profiles were recorded on a LSRFortessa (BD Biosciences) or a FACSymphony A5 instrument (BD Biosciences). Data was analyzed using FlowJo Version 10.7.1. Cell sorting experiments were performed on a BD FACSAria Fusion instrument. Post-sort quality control was performed for each sample.

### RNA extraction, cDNA synthesis and qPCR

Total RNA was isolated using Isolate II RNA Mini Kit (Bioline). cDNA was synthesized using SensiFAST cDNA Synthesis Kit (Bioline). Quantitative real time PCR was performed with iTaq Universal SYBR Green Supermix (BioRad) in a CFX96 Touch BioRad system. Expression levels were normalized to GAPDH. For a primer list see Extended Table 1.

### CUT&RUN protocol

The CUT&RUN protocol was slightly modified as described^37,38^. In brief, trypsinized or cell sorted ESC (between 150,000-500,000 cells depending on the experiment) were washed three times with Wash Buffer (20 mM HEPES-KOH pH 7.5, 150 mM NaCl, 0.5 mM spermidine, Roche complete Protease Inhibitor tablet EDTA free) and bound to activated Concanavalin A beads (Polysciences) for 10 minutes at room temperature. Cells were then permeabilized in Digitonin Buffer (0.05 % Digitonin and 0.1% BSA in Wash Buffer) and incubated with the antibody against CTCF (07-729, Millipore) at 4°C for 2 hours. For negative controls, Guinea Pig anti-Rabbit IgG (ABIN101961, Antibodies-online) was used. Cells were washed with Digitonin Buffer following antibody incubation, and further incubated with purified hybrid protein A-protein G-Micrococcal nuclease (pAG-MNase) at 4°C for 1 hour. Samples were washed in Digitonin Buffer, resuspended in 150 μl Digitonin Buffer and equilibrated to 0°C on ice water for 5 minutes. To initiate MNase cleavage, 3 μl 100 mM CaCl_2_ was added to cells and after 1 hour of digestion, reactions were stopped with the addition of 150 μl 2x Stop Buffer (340 mM NaCl, 20 mM EDTA, 4 mM EGTA, 0.02 % Digitonin, 50 μg/ml RNase A, 50 μg/ml Glycogen). Samples were incubated at 37°C for 10 minutes to release DNA fragments and centrifuged at 16,000 g for 5 minutes. Supernatants were collected and a mix of 1.5 μl 20% SDS / 2.25 μl 20 mg/ml Proteinase K was added to each sample and incubated at 65°C for 35 minutes. DNA was precipitated with ethanol and sodium acetate and pelleted by high-speed centrifugation at 4°C, washed, air-dried and resuspended in 10 μ 0.1x TE.

### Library preparation and sequencing

The entire precipitated DNA obtained from CUT&RUN was used to prepare Illumina compatible sequencing libraries. In brief, end-repair was performed in 50 μl of T4 ligase reaction buffer, 0.4 mM dNTPs, 3 U of T4 DNA polymerase (NEB), 9 U of T4 Polynucleotide Kinase (NEB) and 1 U of Klenow fragment (NEB) at 20°C for 30 minutes. End-repair reaction was cleaned using AMPure XP beads (Beckman Coulter) and eluted in 16.5 μl of Elution Buffer (10 mM Tris-HCl pH 8.5) followed by A-tailing reaction in 20 μl of dA-Tailing reaction buffer (NEB) with 2.5 U of Klenow fragment exo- (NEB) at 37°C for 30 minutes. The 20 μl of the A-tailing reaction were mixed with Quick Ligase buffer 2X (NEB), 3000 U of Quick Ligase (NEB) and 10 nM of annealed adaptor (Illumina truncated adaptor) in a volume of 50 μl and incubated at room temperature for 20 min. The adaptor was prepared by annealing the following HPLC-purified oligos: 5’-Phos/GATCGGAAGAGCACACGTCT-3’ and 5’-ACACTCTTTCCCTACACGACGCTCTTCCGATC*T-3’ (*phosphorothioate bond). Ligation was stopped by adding 50 mM of EDTA, cleaned with AMPure XP beads and eluted in 14 μl of Elution Buffer. All volume was used for PCR amplification in a 50 μl reaction with 1 μM primers TruSeq barcoded primer p7, 5’-CAAGCAGAAGACGGCATACGAGATXXXXXXXXGTGACTGGAGTTCAGACGTGTGCTCTTCCGATC*T-3’ and TruSeq barcoded primer p5 5’-AATGATACGGCGACCACCGAGATCTACACXXXXXXXXACACTCTTTCCCTACACGACGCTCTTCCGATC*T-3’ (* represents a phosphothiorate bond and XXXXXXXX a barcode index sequence), and 2X Kapa HiFi HotStart Ready mix (Kapa Biosciences). The temperature settings during the PCR amplification were 45 s at 98°C followed by 15 cycles of 15 s at 98°C, 30 s at 63°C, 30 s at 72°C and a final 5 min extension at 72°C. PCR reactions were cleaned with AMPure XP beads (Beckman Coulter), run on a 2% agarose gel and a band of 300bp approximately was cut and gel purified using QIAquick Gel Extraction Kit (QIAGEN). Library concentration was determined with KAPA Library Quantification Kit for Illumina Platforms (Kapa Biosystems). Sequencing was performed on the Illumina NextSeq550 (75bp pair-end reads).

### Cut&Run data processing

Data were processed using a modified version of Cut&RunTools^39^. Reads were adapter trimmed using fastp v.0.20.0^40^. An additional trimming step was performed to remove up to 6bp adapter from each read. Next, reads were aligned to the mm10 genome using bowtie2^40^ with the ‘dovetail’ and ‘sensitive’ settings enabled. Alignments were further divided into ≤ 120-bp and > 120-bp fractions. macs2^41^ was used to call peaks with q-value cutoff < 0.01. Normalized (RPKM) signal tracks were generated using the ‘bamCoverage’ utility from deepTools with parameters bin-size=25, smooth length=75, and ‘center_reads’ and ‘extend_reads’ options enabled^42^.

### Processing for published ChIP datasets

Reads were aligned to the mm10 genome using bowtie2^40^. Duplicate reads were removed using MarkDuplicates from the Picard toolkit (“Picard Toolkit.” 2019. Broad Institute, GitHub Repository. http://broadinstitute.github.io/picard/). Normalized (RPKM) signal tracks were generated bamCoverage utility from deepTools^43^, using the parameters bin-size=25, smooth length=75, ‘center_reads’ and ‘extend_reads’. For paired-end data, read mates were extended to the fragment size defined by the two read mates. For single-end ChIP-seq data, reads were extended to the estimated fragment length estimated by phantompeakqualtools^44^.

### RNAseq data processing and batch correction

Fastq files for RNAseq experiments ^5,23^ were downloaded from SRA. RNAseq reads were adapter trimmed using fastp v.0.20.0 (Chen et al., 2018). Transcript expression was quantified via mapping to mouse gencode v25 transcripts using salmon (Patro et al., 2017). In order to compare the two RNAseq experiments, batch correction was performed. Gene counts across samples were quantile-normalized using the limma package^44^. Batch correction was then performed on quantile-normalized counts using COMBAT^45^. Gene association was performed by using GREAT (http://great.stanford.edu/public/html/) using “single nearest gene” by default 1000kb distance.

### ENDseq

END-seq was performed as described^47^. Briefly, for untreated DOX-treated ESC^DUX^, a total of 30 million cells in single cell suspension were embedded in a single agarose plug. Lysis and digestion of embedded cells was performed using Proteinase K (50°C, 1 hour then 37°C for 7 hours). Agarose plugs were rinsed in TE buffer and treated with RNase A at 37°C, 1 hour. Next, DNA ends were blunted. For these reactions, DNA was retained in the plugs to prevent shearing. The first blunting reaction was performed using ExoVII (NEB, M0379S) for 1hr, 37C. Plugs were washed twice in NEB Buffer 4 (1X), immediately followed by the second blunting reaction using ExoT (NEB, M0265S) for 1 hour, 24°C. After this final blunting, two washes were performed in NEBNext dA-Tailing Reaction Buffer (NEB, B6059S), followed by A-tailing (Klenow 3’-> 5’ exo-, NEB, M0212S). After A-tailing, we performed a ligation with the “END-seq hairpin adaptor 1,” listed in reagents section, using NEB Quick Ligation Kit (NEB, M2200S).

### DNA sonication, End-Repair, A-tailing, and Library Amplification

Agarose plugs were then melted and dissolved. DNA was sonicated using to a median shear length of 170bp using a Covaris S220 sonicator for 4 min at 10% duty cycle, peak incident power 175, 200 cycles per burst, 4°C. Following the sonication, DNA was precipitated with ethanol and dissolved in 70 μl TE buffer. 35 μL of Dynabeads were washed twice with 1 mL Binding and Wash Buffer (1xBWB) (10 mM Tris-HCl pH8.0, 1 mM EDTA, 1 M NaCl, 0.1% Tween20). After the wash, beads were recovered using a DynaMag-2 magnetic separator (12321D, Invitrogen) and supernatants were discarded. Washed beads were resuspended in 130 μL 2xBWB (10 mM Tris-HCl pH8.0, 2 mM EDTA, 2 M NaCl) combined with the 130 μL of sonicated DNA followed by an incubation at 24°C for 30 min. Next, the supernatant was removed, and the biotinylated DNA bound to the beads was washed thrice with 1 mL 1xBWB, twice with 1 mL EB buffer, once with 1 mL T4 ligase reaction buffer (NEB) and then resuspended in 50 μL of end-repair reaction mix (0.4 mM of dNTPs, 2.7 U of T4 DNA polymerase (NEB), 9 U of T4 Polynucleotide Kinase (NEB) and 1 U of Klenow fragment (NEB)) and incubated at 24°C for 30 min. Once again, the supernatant was removed using a magnetic separator and beads were then washed once with 1 mL 1xBWB, twice with 1 mL EB buffer, once with 1 mL NEBNext dA-Tailing reaction buffer (NEB) and then resuspended in 50 μL of with NEBNext dA-Tailing reaction buffer (NEB) and 20 U of Klenow fragment exo- (NEB). The A-tailing reaction was incubated at 37°C for 30 min. The supernatant was removed using a magnetic separator and washed once with 1 mL NEBuffer 2 and resuspended in 115 mL of Ligation reaction with Quick Ligase buffer (NEB), 6,000 U of Quick Ligase (NEB) and ligated to “END-seq hairpin adaptor 2” by incubating the reaction at 25°C for 30 min. Reaction was stopped by adding 50 mM of EDTA, and beads washed 3X BWB, 3X EB, and eluted in 8 μL of EB. Hairpin adaptors were digested using USER enzyme (NEB, M5505S) at 37C for 30 minutes. PCR amplification was performed in 50 μL reaction with 10 mM primers 5’-CAAGCAGAAGACGGCATACGA-GATXXXXXXGTGACTGGAGTTCAGACGTGTGCTCTTCCGATC*T-3’ and 5’-AATGATACGGCGACCACCGAGATCTACACTCTTTCCCTACACGACGCTCTTCCGATC*T-3’, and 2X Kapa HiFi HotStart Ready mix (Kapa Biosciences). * represents a phosphothioratebond and NNNNNN a Truseq index sequence. PCR program: 98°C, 45 s; 15 cycles [98°C, 15 s; 63°C, 30 s; 72°C, 30 s]; 72°C, 5 min. PCR reactions were cleaned with AMPure XP beads, and after running the reactions on a 2% agarose gel, 200-500 bp fragments were isolated. Libraries were purified using QIA-quick Gel Extraction Kit (QIAGEN). Library concentration was determined with KAPA Library Quantification Kit for Illumina Platforms (Kapa Biosystems) and the sequencing was performed on Illumina NextSeq 500 or 550 (75bp single end reads).

### Processing of ENDseq data

END-seq reads were aligned to the mouse reference genome mm10 using bowtie (v1.1.2)^41^ (PMID: 19261174) with parameters -n 3 -l 50 -k 1. Functions “view” and “sort” of samtools (v 1.6) (PMID: 19505943) were used to convert and sort the aligned sam files to sorted bam files. Bam files were further converted to bed files by bedtools bamToBed command (PMID: 20110278). END-seq peaks were called by MACS (v1.4.3)^42^ with parameters --nolambda --nomodel --keep-dup=all (PMID: 18798982) and peaks within blacklisted regions (https://sites.google.com/site/anshulkundaje/projects/blacklists) were filtered out (PMID: 31249361). Overlapped peaks from two independent clones were used in this paper.

## ACKNOWLEDGMENTS

We thank Bechara Saykali and Pedro Rocha for critical reading of the manuscript, and to Jacob Paiano for critical discussion. We are grateful to Christian Franke for the continuous technical support on R. We also thank Pedro Rocha, Rafael Casellas and Seol Kyoung Jung for their help on exploring HiC data. David Goldstein and the CCR Genomics Core for sequencing support and Ferenc Livak and the CCR Flow cytometry Core for experimental support. Research in S.R. laboratory is supported by the Intramural Research Program of the NIH. T.O. is supported by a postdoctoral fellowship of the Helen Hay Whitney Foundation.

## AUTHOR CONTRIBUTIONS

T.O. and S.R. conceived the study. T.O., M.V-S. designed, performed and analyzed experiments. C.N.D. and M.F. provided technical support. D.T. and P.C.F. analyzed sequencing data. G.P. provided support with high-throughput microscopy imaging. A.D.T. and M.J.K. analyzed confocal microscopy data. E.L.D. and M.M.P. provided critical reagents. N.Z. performed ENDseq experiments. W.W. analyzed ENDseq data. A.N. supervised ENDseq experiments. E.P.N. provided critical reagents. S.R. supervised the study and wrote the manuscript with comments and help from all authors.

## SUPPLEMENTARY INFORMATION

Supplementary Information is available for this paper.

