## Supplementary Information for "CTCF is a Barrier for Totipotent-like Reprogramming"

[1] Laboratory of Genome Integrity, [2] Genetics Branch, [3] Laboratory of Cancer Biology and Genetics, [4] Laboratory of Receptor Biology and Gene Expression and [5] Genome Analysis Unit, CCR, NCI, NIH, Bethesda, MD, USA. [5]Cardiovascular Research Institute, University of California San Francisco, San Francisco, CA, 94143, USA. [6]Department of Biochemistry and Biophysics, University of California San Francisco, San Francisco, CA, 94143, USA.

[5] Lead Contact

### Figure S1

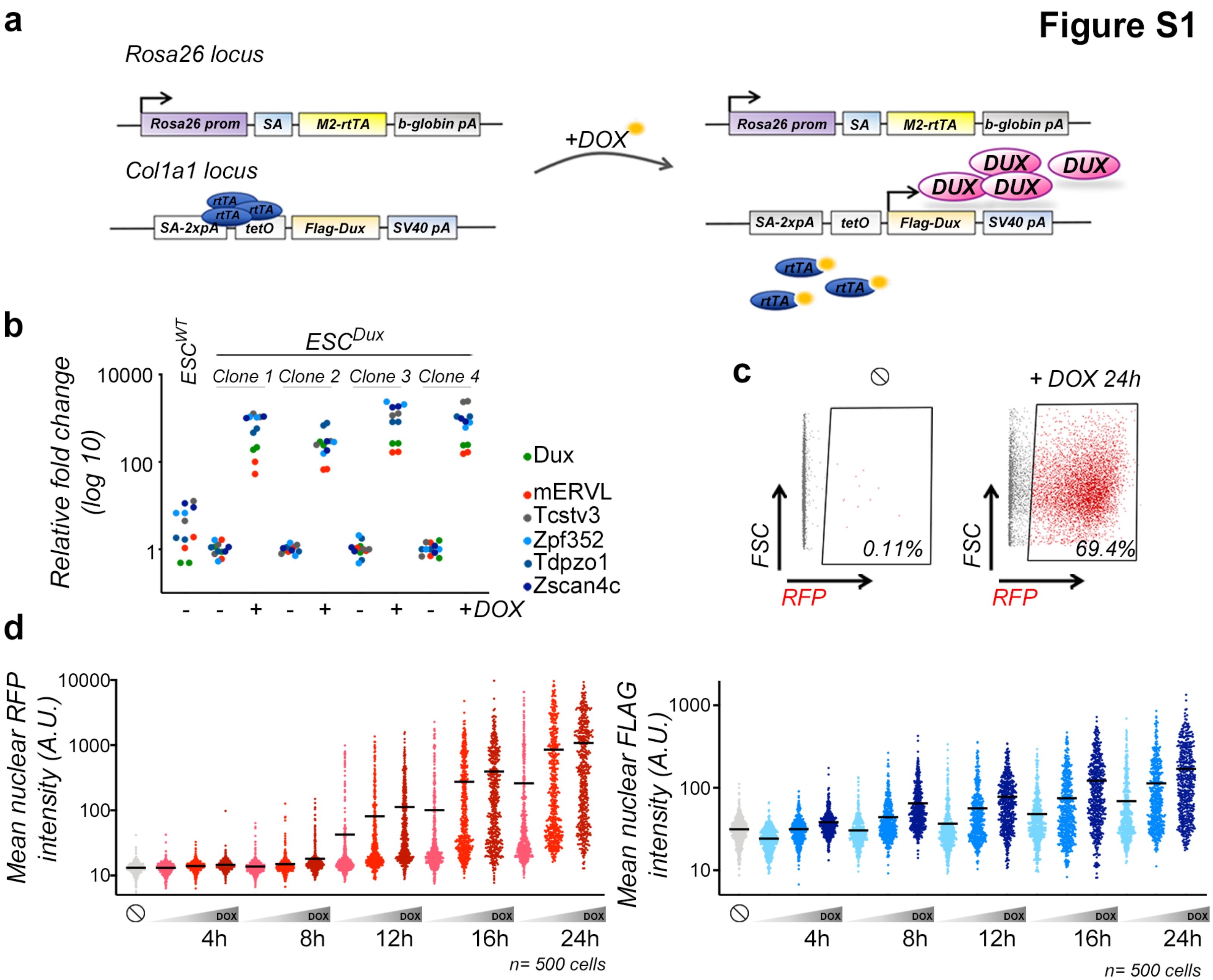
a
Rosa26 locus
Rosa26 prom
SA
M2-rtTA
b-globin pA
Col1a1 locus
rtTA
SA-2xpA
tetO
Flag-Dux
SV40 pA
+DOX
DUX
b
ESC^WT
ESC^Dux
Clone 1
Clone 2
Clone 3
Clone 4
Relative fold change (log 10)
10000
100
1
Dux
mERVL
Tcstv3
Zpf352
Tdpzo1
Zscan4c
- - + - + - + - +DOX
c
+ DOX 24h
FSC
RFP
0.11%
69.4%
d
Mean nuclear RFP intensity (A.U.)
10000
1000
100
10
DOX
4h
8h
12h
16h
24h
n= 500 cells
Mean nuclear FLAG intensity (A.U.)
1000
100
10
DOX
4h
8h
12h
16h
24h
n= 500 cells

**Extended data figure 1: ESC$^{DUX}$ recapitulated all 2C-like features upon DOX-induction. a,** Schematic representation of the two-allele system used to generate ESC$^{DUX}$. DUX cDNA was placed under the control of a tet-responsible sequence (tetO) at the *Col1a1* locus while rtTA is expressed from the *Rosa26* locus[11]. **b,** Graph showing the relative fold change (log10) expression of six 2C associated genes in ESC$^{DUX}$ untreated or treated with DOX for 24 hours. Four ESC clones were tested but two were selected for further downstream analyses. Real-time PCR reactions were performed by triplicate and one representative experiment is shown. GAPDH expression was used to normalize gene expression. **c,** Flow cytometry analysis performed in untreated or DOX-treated *LTR-RFP* reporter ESC$^{DUX}$ for 24 hours. Percentages of RFP$^{+}$ are included within the plots. ∅=No treatment. **d,** HTI quantification of FLAG (left plot) and RFP (right plot) expression in untreated or dox-treated for the indicated times with different DOX concentrations in *LTR-RFP* reporter ESC$^{DUX}$. Center lines indicate mean values. In (**c**, **d**), one representative experiment is shown but at least two independent experiments using multiple clones were performed.

#### Figure S2

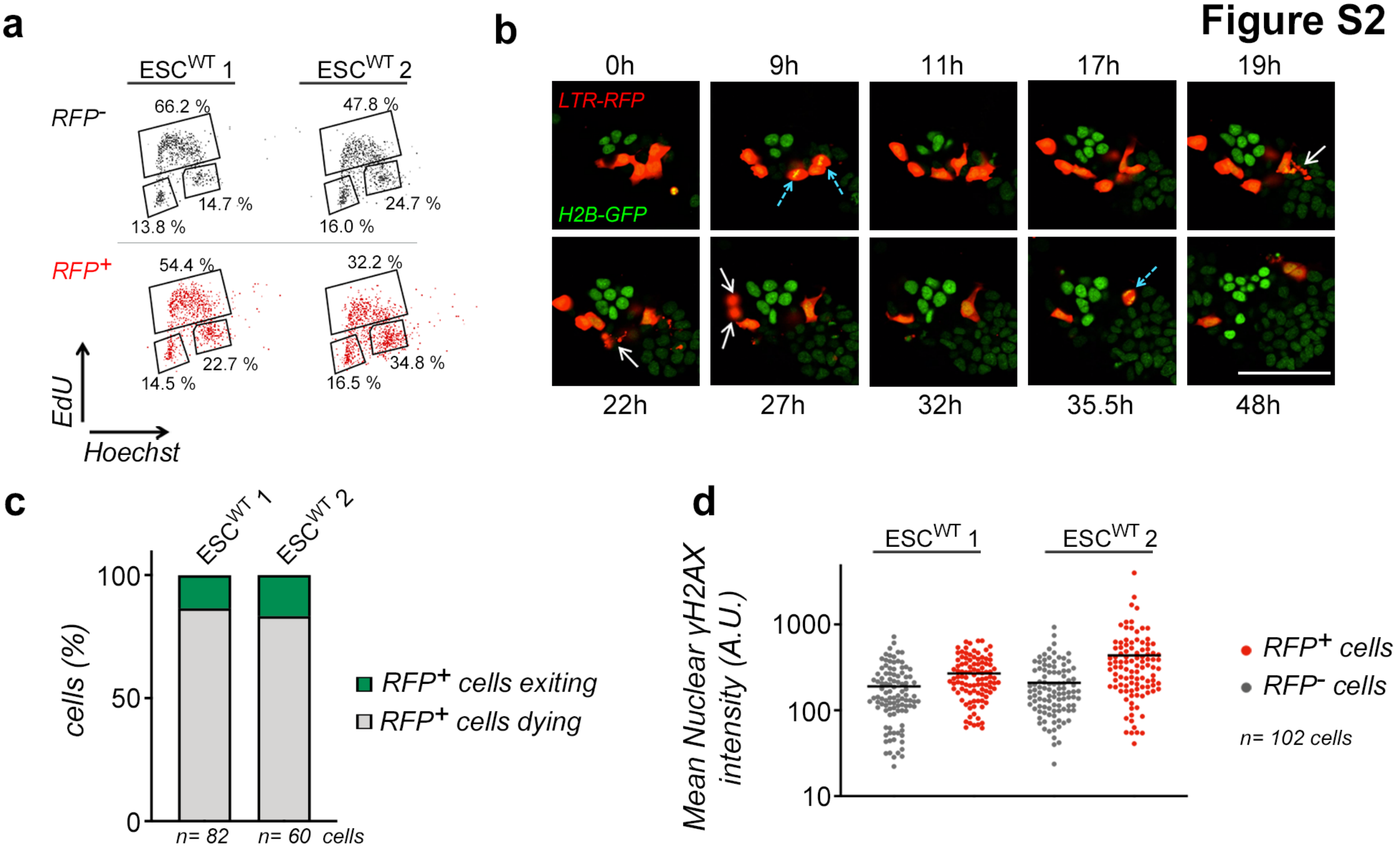
a
ESC^WT 1
ESC^WT 2
RFP-
RFP+
66.2 %
14.7 %
13.8 %
47.8 %
24.7 %
16.0 %
54.4 %
22.7 %
14.5 %
32.2 %
34.8 %
16.5 %
EdU
Hoechst
b
0h
9h
11h
17h
19h
LTR-RFP
H2B-GFP
22h
27h
32h
35.5h
48h
c
ESC^WT 1
ESC^WT 2
100
50
0
cells (%)
RFP+ cells exiting
RFP+ cells dying
n= 82
n= 60 cells
d
ESC^WT 1
ESC^WT 2
Mean Nuclear γH2AX intensity (A.U.)
1000
100
10
RFP+ cells
RFP- cells
n= 102 cells

**Extended data figure 2: Endogenous 2C-like cells showed G2 arrest and increased γH2AX. a,** Flow cytometry analysis of the cell cycle distribution in two wild type *LTR-RFP* reporter ESC lines (R1 and G4 ESC). Cells were split in $RFP^-$ or $RFP^+$ subpopulations and percentages for each phase of the cells cycle are included. One representative experiment is shown but at least two independent experiments were performed. **b,** Representative images obtained from a live cell time lapse experiment where *LTR-RFP* reporter R1 ESC expressing H2B-eGFP were followed over time. Indicated times showed the time when recording started. Two independent wild type *LTR-RFP* reporter ESC lines (R1 and G4) were imaged in two independent experiments but only one representative ESC line (R1) is shown. White arrows showed $RFP^+$ ESC undergoing cell death. Blue dashed arrows showed $RFP^+$ ESC undergoing cell division. Scale bar, 100 μm. **c,** Plot showing the percentage of endogenous cycling 2C-like cells in two wild-type ESC lines that successfully exit from the totipotent-like state and return to pluripotency or undergo cell death. Quantification was performed by following the fate of a total of 82 and 60 2C-like cells per cell line in the time-lapse experiments from (**b**). **d,** HTI quantification of γH2AX in two wild type *LTR-RFP* reporter ESC lines (R1 and G4 ESC). Cells were split in $RFP^-$ and $RFP^+$ subpopulations. Center lines indicate mean values. One representative experiment is shown but at least two independent experiments were performed.

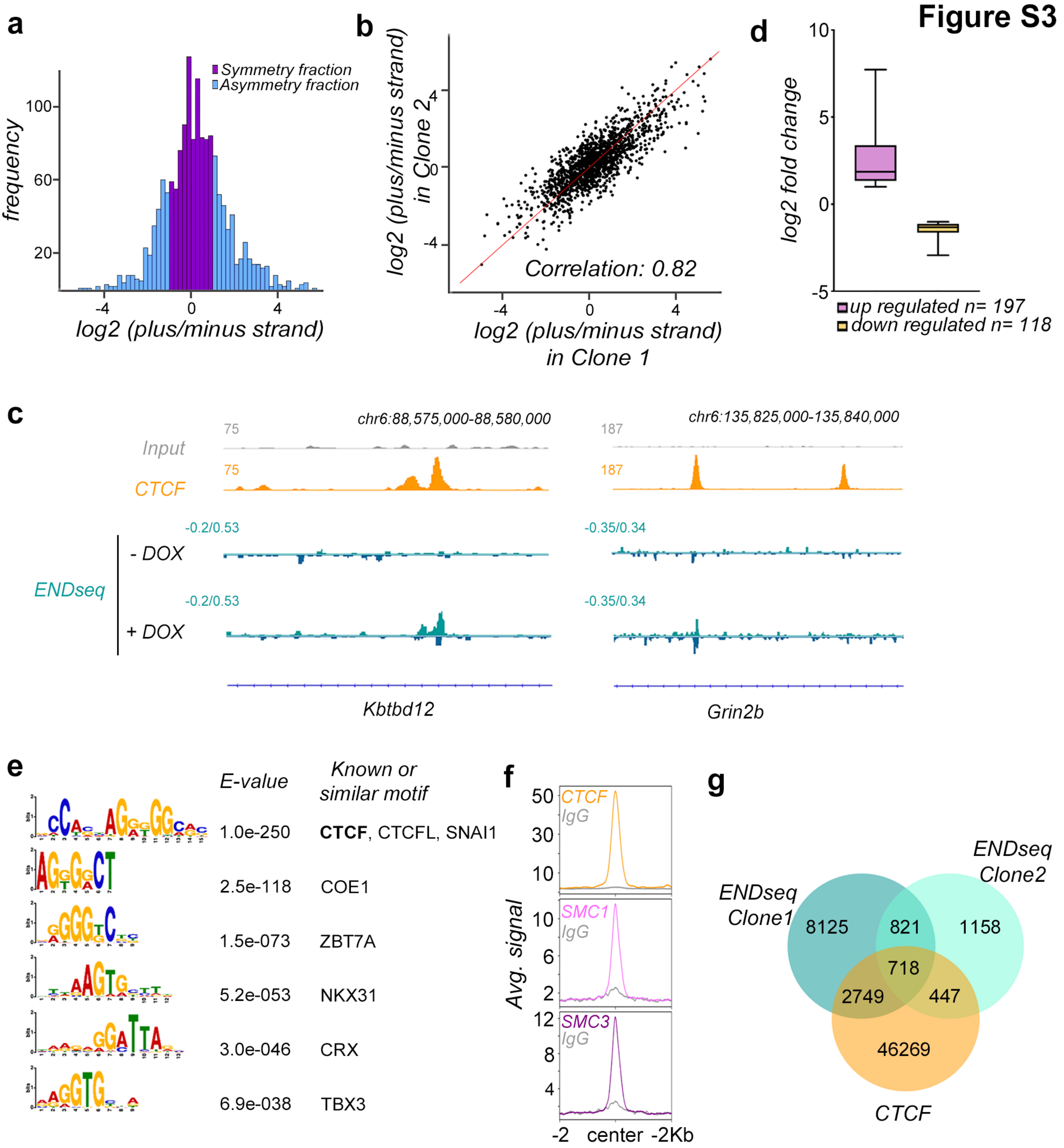
Figure S3
a
frequency
100
60
20
-4
0
4
log2 (plus/minus strand)
Symmetry fraction
Asymmetry fraction
b
log2 (plus/minus strand) in Clone 2
log2 (plus/minus strand) in Clone 1
Correlation: 0.82
d
log2 fold change
10
5
0
-5
up regulated n= 197
down regulated n= 118
c
chr6:88,575,000-88,580,000
chr6:135,825,000-135,840,000
Input
CTCF
75
187
-0.2/0.53
-0.35/0.34
- DOX
+ DOX
ENDseq
Kbtbd12
Grin2b
e
E-value
Known or similar motif
1.0e-250
CTCF, CTCFL, SNAI1
2.5e-118
COE1
1.5e-073
ZBT7A
5.2e-053
NKX31
3.0e-046
CRX
6.9e-038
TBX3
f
Avg. signal
CTCF
IgG
SMC1
SMC3
-2
center
-2Kb
g
ENDseq Clone1
ENDseq Clone2
CTCF
8125
821
1158
718
2749
447
46269

**Extended data figure 3: CTCF is enriched in ENDseq sites.**

**a,** Histogram plot showing the frequency of single (asymmetric fraction) or double (symmetric fraction) strand breaks observed within the ENDseq peaks. Ratio between the signal intensity per strand was used to define symmetric or asymmetric ENDseq peaks. **b,** Plot showing the level of correlation between the type of DNA lesions (single or double strand breaks) observed in two independent $ESC^{DUX}$ clones. Pearson correlation value is shown. **c,** Genome browser tracks showing ENDseq signal in untreated and DOX-treated $ESC^{DUX}$ at the indicated genome location. One representative single (left panel) or double (right panel) end break is shown. Input (IgG) is shown as a background reference control. **d,** Graph showing the relative fold change (log2) in genes differentially expressed (a total of 314 genes) between wild-type and DUX-expressing ESC located near ENDseq sites (Supplementary Table 5). Data was obtained from[5]. **e,** Results from the motif enrichment analysis performed on the set of overlapping 1539 ENDseq sites identified in DUX-expressing ESC. **f,** Plots showing CTCF[20], SMC1 and SMC3[21] enrichment at the set of 1539 ENDseq sites identified in DOX-treated $ESC^{DUX}$. Input (IgG) is also shown as a background reference control. **g,** Venn diagram showing the number of ENDseq and CTCF overlapping peaks between the two $ESC^{DUX}$ clones analyzed. Peak calling for CTCF binding sites in ESC was obtained from[20].

### Figure S4

a

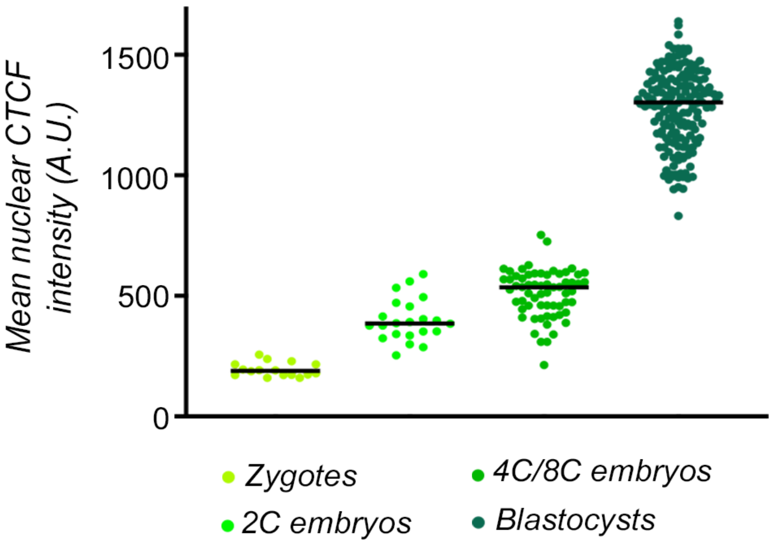
Mean nuclear CTCF intensity (A.U.)
1500
1000
500
0
Zygotes
2C embryos
4C/8C embryos
Blastocysts

b

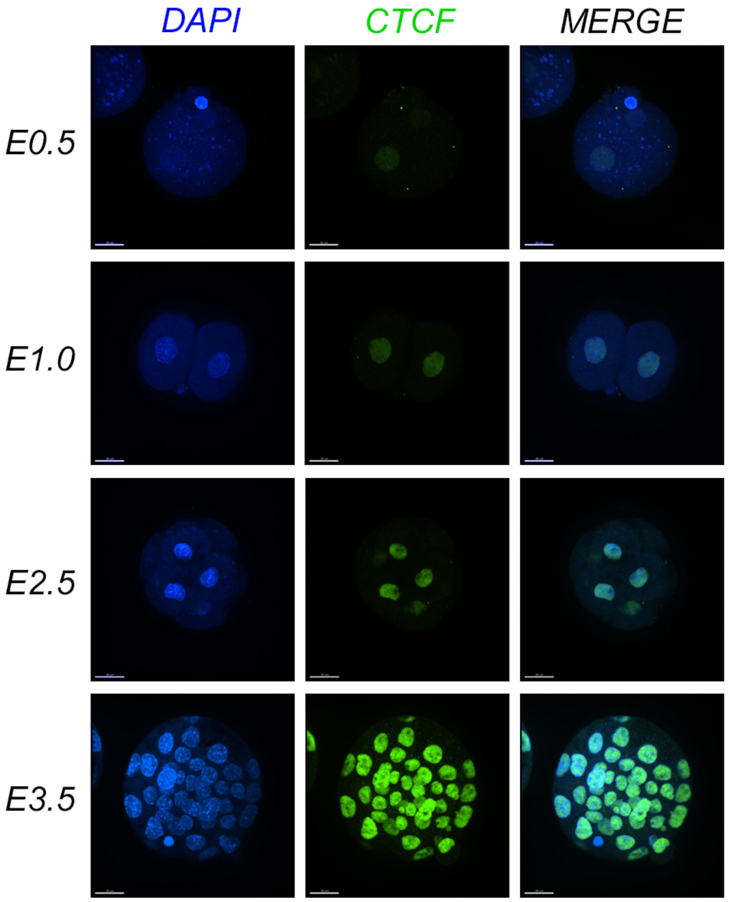
DAPI
CTCF
MERGE
E0.5
E1.0
E2.5
E3.5

**Extended data figure 4: CTCF expression levels increased during embryonic development; a,** Graph showing CTCF mean nuclear intensity in mouse embryos at different developmental stages, Zygotes (E0.5, 8 embryos), 2C (E1.0, 11 embryos), 4C/8C (E2.5, 14 embryos) and blastocysts (E3.5, 7 embryos). **b,** Representative images from the mouse embryos described in (**a**). Scale bar, 20μm.

### Figure S5

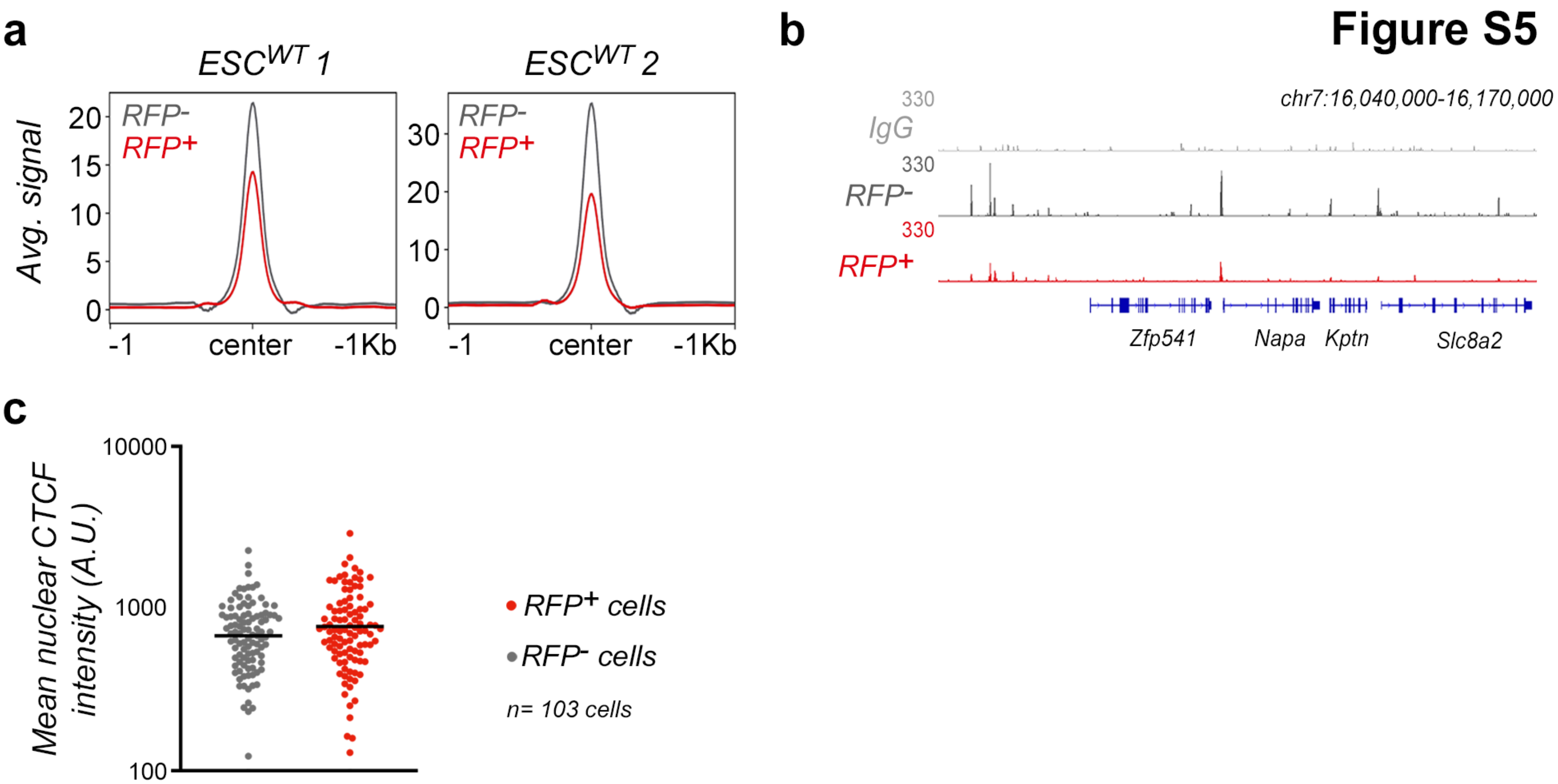
a
ESC^WT 1
ESC^WT 2
Avg. signal
RFP-
RFP+
20
15
10
5
0
30
20
10
0
-1
center
-1Kb
b
chr7:16,040,000-16,170,000
330
IgG
330
RFP-
330
RFP+
Zfp541
Napa
Kptn
Slc8a2
c
Mean nuclear CTCF intensity (A.U.)
10000
1000
100
RFP+ cells
RFP- cells
n= 103 cells

**Extended data figure 5: Endogenous 2C-like cells showed decreased chromatin bound CTCF. a,** Cut&Run read density plot (RPKM) showing CTCF occupancy in the set of ESC-specific 50183 CTCF sites in $RFP^+$ and $RFP^-$ sorted ESC obtained from *LTR-RFP* reporter R1 and E14 ESC. The signal obtained in corresponding inputs (IgG) is subtracted. **b,** Genome browser tracks showing CTCF occupancy at the indicated genome location from samples described in (**a**). Input (IgG) is also shown as a reference control. **c,** HTI quantification of CTCF in wild type *LTR-RFP* reporter ESC line. Cells were split in $RFP^+$ or $RFP^-$ subpopulations. Center lines indicate mean values. Representative data from one wild type *LTR-RFP* reporter ESC line is shown but two ESC lines were assayed.

### Figure S6

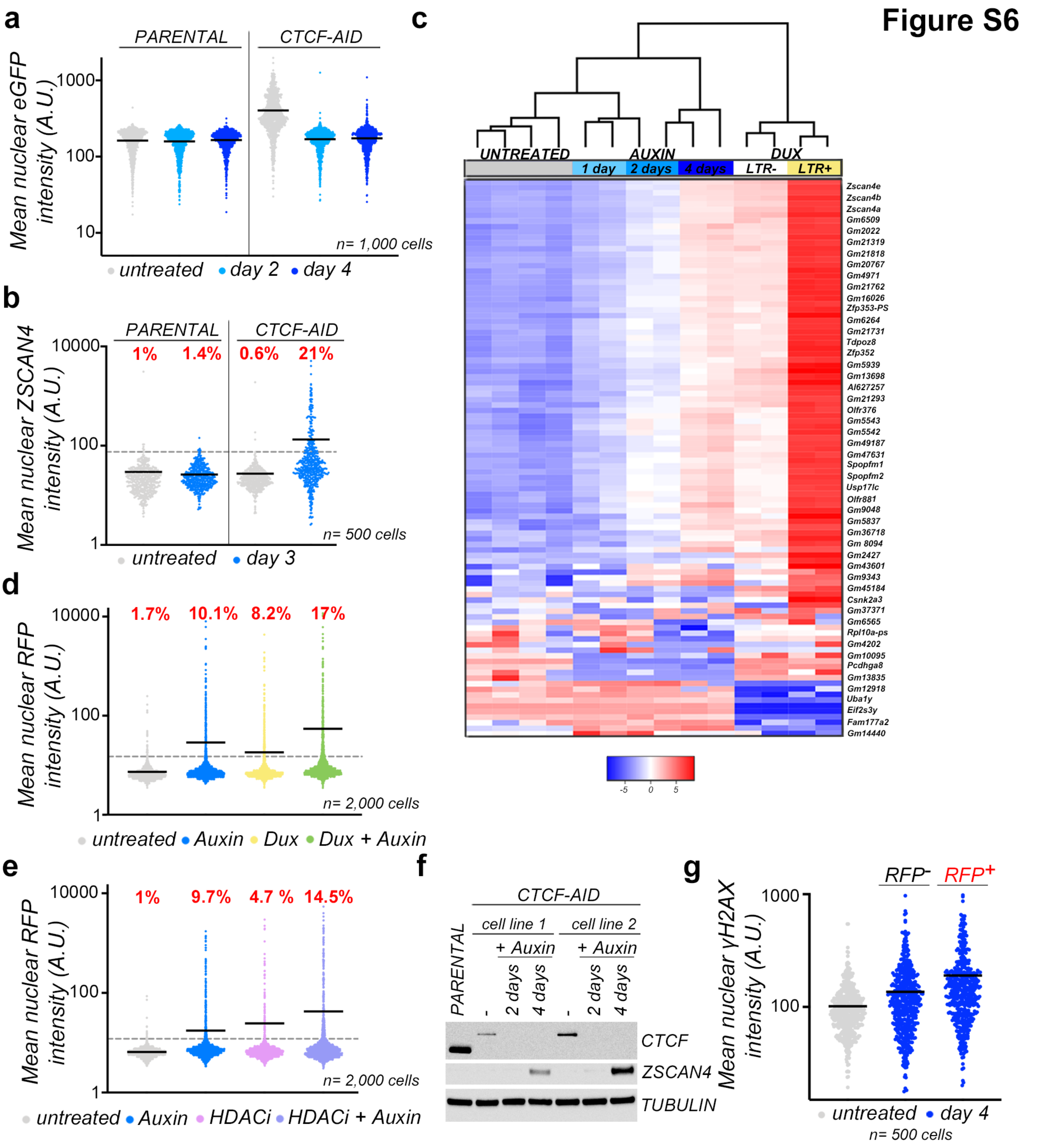
a
PARENTAL
CTCF-AID
Mean nuclear eGFP intensity (A.U.)
1000
100
10
n= 1,000 cells
untreated
day 2
day 4
b
PARENTAL
CTCF-AID
1%
1.4%
0.6%
21%
Mean nuclear ZSCAN4 intensity (A.U.)
10000
100
1
n= 500 cells
untreated
day 3
c
UNTREATED
AUXIN
DUX
1 day
2 days
4 days
LTR-
LTR+
Zscan4e
Zscan4b
Zscan4a
Gm6509
Gm2022
Gm21319
Gm21818
Gm20767
Gm4971
Gm21762
Gm16026
Zfp353-PS
Gm6264
Gm21731
Tdpoz8
Zfp352
Gm5939
Gm13698
AI627257
Gm21293
Olfr376
Gm5543
Gm5542
Gm49187
Gm47631
Spopfm1
Spopfm2
Usp17lc
Olfr881
Gm9048
Gm5837
Gm36718
Gm 8094
Gm2427
Gm43601
Gm9343
Gm45184
Csnk2a3
Gm37371
Gm6565
Rpl10a-ps
Gm4202
Gm10095
Pcdhga8
Gm13835
Gm12918
Uba1y
Eif2s3y
Fam177a2
Gm14440
-5
0
5
d
1.7%
10.1%
8.2%
17%
Mean nuclear RFP intensity (A.U.)
10000
100
1
n= 2,000 cells
untreated
Auxin
Dux
Dux + Auxin
e
1%
9.7%
4.7 %
14.5%
Mean nuclear RFP intensity (A.U.)
10000
100
1
n= 2,000 cells
untreated
Auxin
HDACi
HDACi + Auxin
f
CTCF-AID
cell line 1
cell line 2
+ Auxin
+ Auxin
PARENTAL
-
2 days
4 days
-
2 days
4 days
CTCF
ZSCAN4
TUBULIN
g
RFP-
RFP+
Mean nuclear γH2AX intensity (A.U.)
1000
100
untreated
day 4
n= 500 cells

**Extended data figure 6: CTCF depletion induces spontaneous 2C-like conversion in ESC. a,** HTI quantification of eGFP in untreated or auxin-treated for two and four days parental and $ESC^{CTCF-AID}$. Center lines indicate mean values. **b,** HTI quantification of $ZSCAN4^{+}$ cells in untreated or auxin-treated for three days parental and $ESC^{CTCF-AID}$. Center lines indicate mean values. Percentages of $ZSCAN4^{+}$ cells above the threshold are indicated. **c,** Heatmap generated from RNAseq data from untreated or auxin-treated $ESC^{CTCF-AID}$ at different time points[23] together with wild-type or DUX-expressing ESC[5]. DUX-expressing ESC were sorted in $eGFP^{+}$ or $eGFP^{-}$ as a result of the activation of the LTR-eGFP reporter[5]. Heatmap shows the top 100 most differentially expressed genes. **d, e** HTI quantification of $RFP^{+}$ cells in untreated or auxin-treated for four days *LTR-RFP* reporter $ESC^{CTCF-AID}$ expressing Dux (**d**) or treated with a histone deacetylase inhibitor (**e**) where indicated. Center lines indicate mean values. Percentages of $RFP^{+}$ cells above the threshold are indicated. **f,** Western blot analysis of the indicated proteins performed in two newly generated $KH2\text{-}ESC^{CTCF-AID}$ treated with auxin for 2 and 4 days. Parental ESC were used to show the smaller size and higher levels of CTCF. Tubulin levels are shown as a loading control. Two ESC cell lines are shown but 4 cell lines from 2 different genetic backgrounds were generated and confirmed. **g,** HTI quantification of γH2AX in untreated or auxin-treated for four days in *LTR-RFP* reporter $ESC^{CTCF-AID}$. Auxin-treated cells were split based in $RFP^{+}$ and $RFP^{-}$. Center lines indicate mean values. In (**a**), (**b**), (**d**), (**e**) and (**g**), one representative experiment is shown but at least two independent experiments were performed by duplicates.

### Figure S7

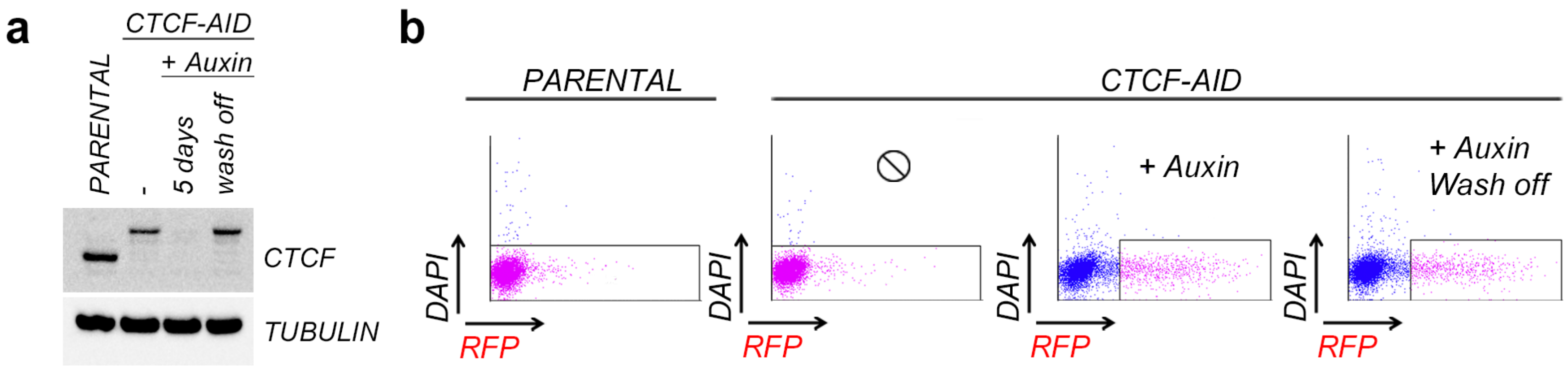
a
CTCF-AID
+ Auxin
PARENTAL
-
5 days
wash off
CTCF
TUBULIN
b
PARENTAL
CTCF-AID
+ Auxin
+ Auxin
Wash off
DAPI
RFP

**Extended data figure 7: Re-expression of CTCF promotes exit from the 2C-like state. a,** Western blot analysis of CTCF performed in untreated or auxin-treated for five days in parental ESC or $ESC^{CTCF\text{-}AID}$. Wash off sample include with four days with auxin plus 18 hours without auxin. Tubulin levels are shown as a loading control. ∅=No treatment. **b,** Flow cytometry plots generated from samples described in (**a**). Gating shows the actual $RFP^{+}$ cells sorted for the experiment.

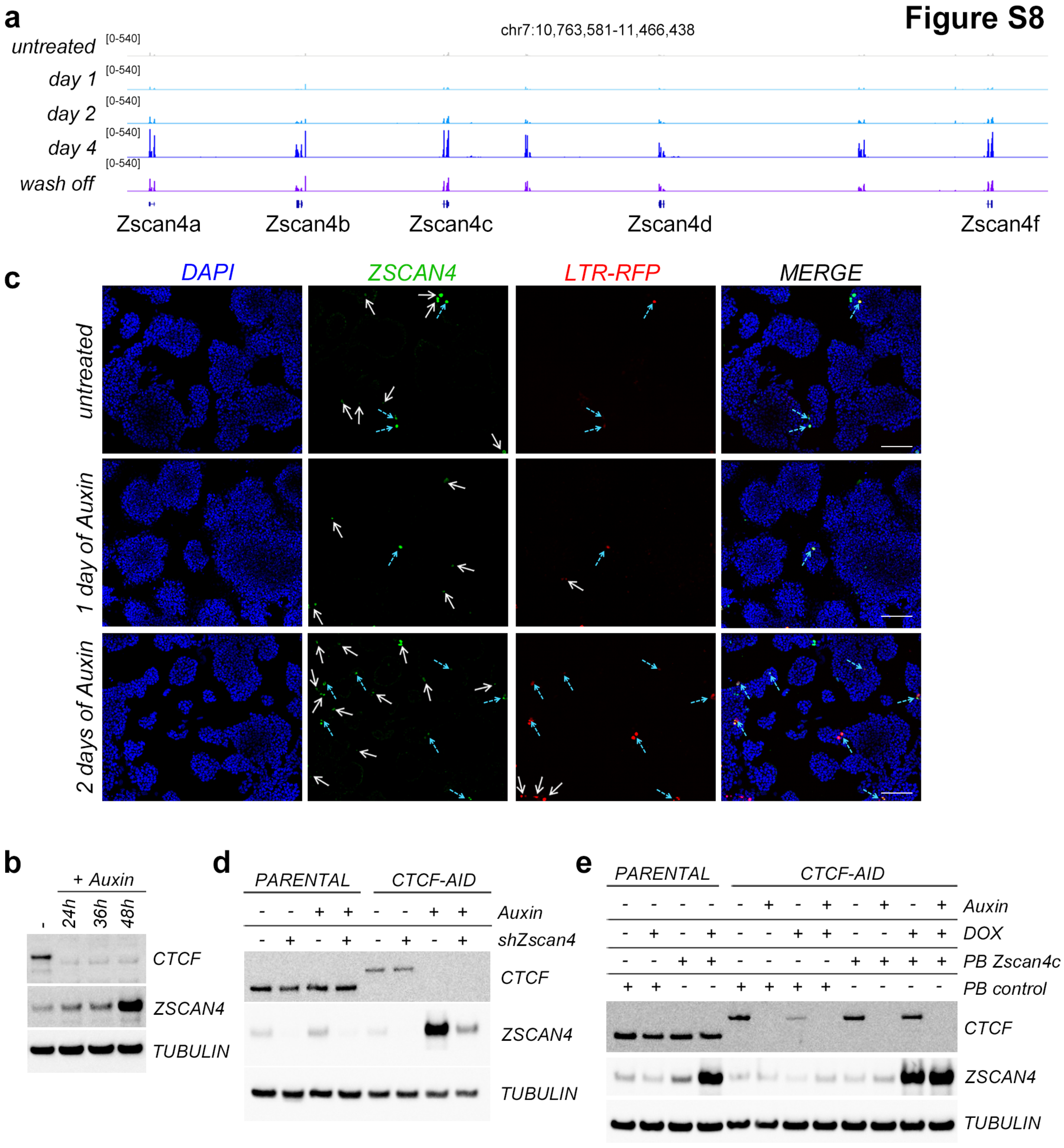

Figure S8
a
chr7:10,763,581-11,466,438
untreated
day 1
day 2
day 4
wash off
[0-540]
Zscan4a
Zscan4b
Zscan4c
Zscan4d
Zscan4f
c
DAPI
ZSCAN4
LTR-RFP
MERGE
untreated
1 day of Auxin
2 days of Auxin
b
+ Auxin
-
24h
36h
48h
CTCF
ZSCAN4
TUBULIN
d
PARENTAL
CTCF-AID
Auxin
shZscan4
CTCF
ZSCAN4
TUBULIN
e
PARENTAL
CTCF-AID
Auxin
DOX
PB Zscan4c
PB control
CTCF
ZSCAN4
TUBULIN

**Extended data figure 8: ZSCAN4 expression is necessary for efficient 2C-like conversion upon CTCF depletion. a,** Genome browser tracks showing RNAseq RPKM read count at the ZSCAN4 cluster in the indicated samples. **b,** Representative immunofluorescence images from $ESC^{CTCF\text{-}AID}$ treated with Auxin for one or two days showing $ZSCAN4^{+}$ cells and $ZSCAN4^{+}$; $LTR\text{-}RFP^{+}$ cells. Scale bar, 200µm. White arrows indicate single $ZSCAN4^{+}$ or $LTR\text{-}RFP^{+}$ cells. Blue dashed arrows indicate double $ZSCAN4^{+}$; $LTR\text{-}RFP^{+}$ cells **c,** Western blot analysis of CTCF and ZSCAN4 performed in untreated or auxin-treated for the indicated hours in $ESC^{CTCF\text{-}AID}$. Tubulin levels are shown as a loading control. **d,** Western blot analysis of CTCF and ZSCAN4 performed in untreated or auxin-treated for four days in parental ESC or $ESC^{CTCF\text{-}AID}$. ESC were infected with lentiviruses expressing siRNAs against ZSCAN4 where indicated. Tubulin levels are shown as a loading control. **e,** Western blot analysis of CTCF and ZSCAN4 performed in untreated or auxin-treated for four days in parental ESC or $ESC^{CTCF\text{-}AID}$. ESC were transfected with PiggyBac (PB) control or expressing ZSCAN4 where indicated. Doxycycline was added to induce PB-induced expression. Tubulin levels are shown as a loading control.

**Extended table 1: List of primers.**

| **Primers (qPCR)** | | |
|---|---|---|
| | forward | reverse |
| Dux | GGAGTGAGAGGCAGATCAGG | CTGCTGACCGAAGTCCAACT |
| Zscan4c | TCTTTCTGGTTGGCAGCTTT | GCCAGGCTTCTGTCAAGAAC |
| Zfp352 | AAGGTCCCACATCTGAAGAA | GGGTATGAGGATTCACCCA |
| Tcstv3 | ACCAGCTGAAACATCCATCC | CCATGGATCCCTGAAGGTAA |
| Sp110 | CACCTGCAAACAAGAAAGCA | AACTCCATGTCCAGGTGAGG |
| Tdpoz1 | GCCCTTGATTTCATTGCCTA | CACTTTCGGCTCCAAGAAAG |
| Dub1 | GCCTTCAAGTGCACAGACAA | GATGTCCAGGAAGCGATCAT |
| Ef1a | AACAGGCGCAGAGGTAAAAA | GCACAGCCTCCTTACACCAT |
| **Primers (Cloning)** | | |
| LTR-NotI-F (Cloning LTR sequences into the PiggyBac construct) | CGATGCGGCCGCGCTGTAGTGGTTATTCCTGGTTGTC | |
| LTR-EcoRI-R (Cloning LTR sequences into the PiggyBac construct) | CGGAGAATTCAAGCTTTGTGCACTGAATCACCT | |
| mDUX-FLAG-MfeI-F (Cloning DUX into pBS31) | GGCATCAATTGACCATGGACTACAAGGATGACGACGATAAGGGCAGCGGAGCTGAGGCTGGCTCTCCAGTGGGAGGAT | |
| mDUX-FLAG-MfeI-R (Cloning DUX into pBS31) | GGCATCAATTGTTACAGCATGTCAAGAAGGGTCTGGTACTC | |
| **Primers (sgRNA)** | forward | reverse |
| pX330_sgRNA-ROSA26-F | CACCGCTCCAGTCTTTCTAGAAGAT | AAACATCTTCTAGAAAGACTGGAGC |
